# Bioinformatic and experimental evidence for suicidal and catalytic plant THI4s

**DOI:** 10.1101/2020.04.29.067249

**Authors:** Jaya Joshi, Guillaume A.W. Beaudoin, Jenelle A. Patterson, Jorge D. García-García, Catherine E. Belisle, Lan-Yen Chang, Lei Li, Owen Duncan, A. Harvey Millar, Andrew D. Hanson

## Abstract

Like fungi and some prokaryotes, plants use a thiazole synthase (THI4) to make the thiazole precursor of thiamin. Fungal THI4s are suicide enzymes that destroy an essential active-site Cys residue to obtain the sulfur atom needed for thiazole formation. In contrast, certain prokaryotic THI4s have no active-site Cys, use sulfide as sulfur donor, and are truly catalytic. The presence of a conserved active-site Cys in plant THI4s and other indirect evidence implies that they are suicidal. To confirm this, we complemented the Arabidopsis *tz-1* mutant, which lacks THI4 activity, with a His-tagged Arabidopsis THI4 construct. LC-MS analysis of tryptic peptides of the THI4 extracted from leaves showed that the active-site Cys was predominantly in desulfurated form, consistent with THI4 having a suicide mechanism *in planta*. Unexpectedly, transcriptome datamining and deep proteome profiling showed that barley, wheat, and oat have both a widely expressed canonical THI4 with an active-site Cys, and a THI4-like paralog (non-Cys THI4) that has no active-site Cys and is the major type of THI4 in developing grains. Transcriptomic evidence also indicated that barley, wheat, and oat grains synthesize thiamin *de novo*, implying that their non-Cys THI4s synthesize thiazole. Structure modeling supported this inference, as did demonstration that non-Cys THI4s have significant capacity to complement thiazole auxotrophy in *Escherichia coli*. There is thus a *prima facie* case that non-Cys cereal THI4s, like their prokaryotic counterparts, are catalytic thiazole synthases. Bioenergetic calculations show that, relative to suicide THI4s, such enzymes could save substantial energy during the grain filling period.

**Abbreviations:** DHAla, dehydroalanine; EST, expressed sequence tag; FDR, false discovery rate; IPTG, β-D-1-thiogalactopyranoside

## INTRODUCTION

The thiazole moiety of thiamin (vitamin B1) is synthesized in yeast (*Saccharomyces cerevisiae*) and other fungi by the thiazole synthase THI4, an unusual suicide enzyme [1,2]. In an iron-dependent reaction, yeast THI4 acquires the sulfur atom for the thiazole ring from a Cys residue in the THI4 active site (Cys 205 in yeast THI4), which leaves behind a dehydroalanine (DHAla) residue and irreversibly inactivates the enzyme (Figure 1) [1,3]. In contrast, THI4s from certain archaea (*Methanococcus igneus* and *M. jannaschii*) are not suicide enzymes but true catalysts that mediate multiple reaction cycles. These THI4s lack an active-site Cys and instead use sulfide (HS^−^) as sulfur donor (Figure 1) [3,4]. It is important to note that *M. igneus* and *M. jannaschii* are hyperthermophilic anaerobes from high-sulfide habitats, and hence that their catalytic THI4s normally operate in, and may require, extreme conditions.

**Figure 1.**
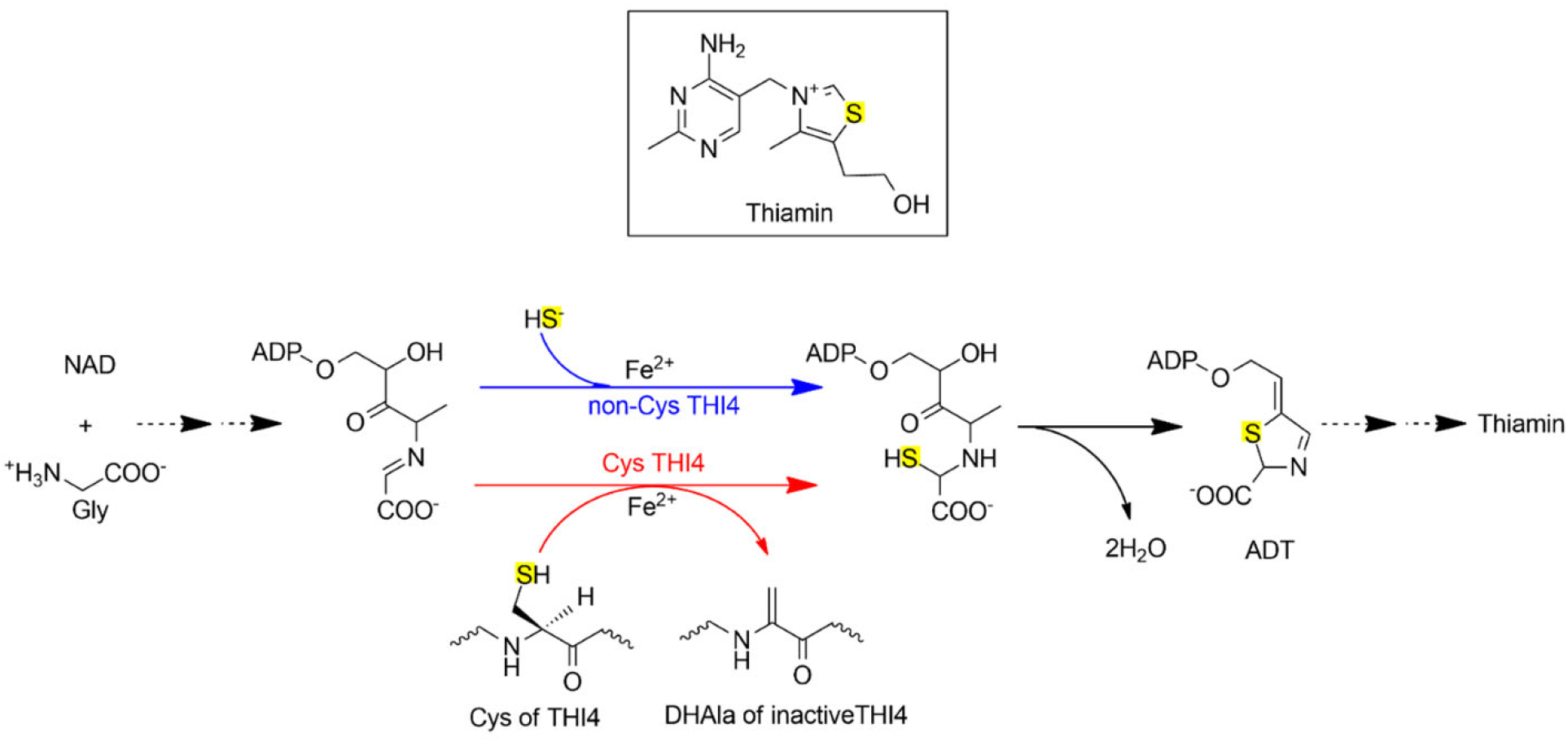
Biosynthesis of the thiazole precursor of thiamin by suicidal and non-suicidal THI4s. THI4s synthesize the adenylated carboxythiazole (ADT) precursor of thiamin from NAD, glycine, and a sulfur atom that in yeast THI4 comes from an active-site Cys residue (Cys 205) and in methanococcal THI4s comes from sulfide (HS^−^). Loss of the Cys sulfur leads to formation of a dehydroalanine (DHAla) residue and irreversible loss of activity. Yeast-type THI4s are consequently suicide enzymes whereas methanococcal THI4s are not suicidal and are true catalysts.

Indirect evidence implies that plant THI4s, like yeast THI4, are suicide enzymes. First, the active-site Cys is conserved in all plant THI4 sequences examined so far [5,6]. Second, the active-site Cys residue in arum lily THI4 was shown to be required for functional complementation of an *Escherichia coli* thiazole auxotroph [6]. Third, when Arabidopsis THI4 was expressed in *E. coli* the active-site Cys was apparently converted to DHAla [1,7]. Fourth, barley [8] and Arabidopsis [9] THI4 proteins turn over faster than all other proteins for which measurements were obtained, as would be expected if THI4 is degraded and replaced after just one reaction cycle [10]. These observations do not, however, prove that plant THI4s operate as suicide enzymes *in planta*. The first aim of this study was accordingly to obtain evidence on this point via peptide mass spectrometry.

The second aim was to investigate whether plant THI4s invariably have an active-site Cys residue or whether some do not and are thus potentially true catalysts. (Henceforth, we term these Cys and non-Cys THI4s, respectively.) This exploration was prompted by the recent findings (i) that non-Cys THI4s occur not only in anaerobic prokaryotes from sulfide-rich environments but also in aerotolerant and microaerophilic – or possibly aerobic – organisms from low-sulfide environments, and (ii) that some of these THI4s can function in *E. coli* in the presence of oxygen [6]. These findings suggested that catalytic THI4s might be able to operate in the conditions found in plant cells, i.e. that no *a priori* biochemical logic bars plants from having catalytic THI4s. When transcriptome and genome database mining and proteomics analysis uncovered non-Cys THI4s in cereal grains, we used comparative genomics, 3D structure modeling, and functional complementation tests in *E. coli* to investigate whether these THI4s are catalytic thiazole synthases. Lastly, we estimated that replacing a suicide THI4 with a catalytic one in cereal grains could substantially increase the ATP supply for starch synthesis.

## MATERIALS AND METHODS

### Chemicals and media

Chemicals and reagents were from Sigma-Aldrich or Fisher Scientific unless otherwise indicated.

### Arabidopsis THI4 expression in the tz-1 mutant

Seed of the *tz-1* mutant [11] was obtained from the Arabidopsis Biological Resource Center (stock number CS3375). Mutant plants were grown in potting soil under fluorescent light (~150 μmol photons m^−2^ s^−1^) in a 12-h-light/12-h-dark cycle at 21-23°C and watered with 100 μM thiamin. The *THI4* promoter and coding sequence were amplified from Arabidopsis Col-0 genomic DNA with primers 1 and 2, while the 3’ sequence was amplified using primers 3 and 4, with primer 3 encoding a His_6_ tag. Primers are listed in Supplementary Table 1. Both amplicons were introduced by TA cloning into pGEM T-Easy (Promega) to generate plasmids pGEM-12 and pGEM-34. pGEM-12 was digested with KpnI and NheI, pGEM-34 with NheI and BamHI and the resulting gene fragments were then introduced into pBIN19 pre-digested with KpnI and BamHI to generate pBIN19-THI4.

pBIN19-THI4 was introduced into *Agrobacterium tumefaciens* strain GV3101 by electroporation and used to transform *tz-1* plants. The resulting transformants were selected on ½ MS media containing 1% sucrose, transferred to soil after three weeks, and allowed to set seed. Seeds from these thiamin prototrophs were then sown in soil. After three weeks, approximately 4 g of seedlings were collected, frozen in liquid nitrogen, and ground to a fine powder. The powder was extracted with 20 ml of buffer A (50 mM sodium phosphate, 300 mM sodium chloride, pH 8.0) for 15 min at 4°C with shaking. The debris was then removed by centrifugation and filtration through Miracloth. One ml of HisPur™ Ni-NTA Superflow Agarose (ThermoFisher Scientific) slurry was added to the extract and incubated for 15 min at 4°C with shaking. The beads were then poured onto a column and washed with 50 ml of buffer A supplemented with 20 mM imidazole. The His_6_-tagged THI4 was eluted with 2 ml of buffer A supplemented with 250 mM imidazole, desalted on a PD-10 column (GE Healthcare) equilibrated with 50 mM ammonium bicarbonate, and lyophilized.

### Arabidopsis THI4 protein analysis

His_6_-tagged THI4 samples were treated with/without alkylation (5 mM dithiothreitol at 56°C for 20 min and then 15 mM iodoacetamide at 22°C for 30 min for Cys alkylation). Alkylated/non-alkylated samples (10 μg) were digested with 0.2 μg trypsin in 50 mM NH_4_HCO_3_ buffer at 37°C overnight. The digest was filtered (0.22 μm) and filtered samples (1-2 μg) were loaded onto a C18 high capacity nano LC chip (Agilent Technologies) using a 1200 series capillary and eluted directly into a 6550 series quadrupole time-of-flight mass spectrometer (Agilent Technologies) with a 1200 series nano pump as described [9]. Agilent .d files were converted to mzML using the Msconvert package (version 2.2.2973) from the Proteowizard project, and mzML files were subsequently converted to Mascot generic files using the mzxml2 search tool from the TPPL version 4.6.2. Mascot generic file peak lists were searched against an in-house Arabidopsis database comprising ATH1.pep (release 10) from The Arabidopsis Information Resource and the Arabidopsis mitochondrial and plastid protein sets (33621 sequences; 13487170 residues) [12], using the Mascot search engine version 2.3 and error tolerances of 100 ppm for MS and 0.5 Da for MS/MS; ‘Max Missed Cleavages’ set to 1; variable modifications of Cys-S, oxidation (Met) and carbamidomethyl (Cys). We used iProphet and ProteinProphet from the Trans Proteomic Pipeline (TPP) to analyze peptide and protein probability and global false discovery rate (FDR) [13-15]. Two separate mass spectrometry runs were combined. The reported peptide lists with p=0.8 have FDRs <3%. Alkylated and non-alkylated peptides were used to identify Cys modifications; non-alkylated peptides were used to identify native N-termini with a semi-trypsin search parameter in Mascot.

### Wheat and barley THI4 proteomics

*Triticum aestivum* cv. Wyalkatchem and *Hordeum vulgare* cv. Morex were grown in soil in long day conditions (450 μmol photons m^−2^ s^−1^ for 16 h at 26°C/8 h dark at 22°C, 60% relative humidity). Endosperm was harvested from imbibed mature seed in wheat and whole developing grains (late dough stage) in barley. Total protein was extracted by grinding in liquid nitrogen followed by methanol/chloroform extraction. Proteins were trypsin-digested and peptides prefractionated into 96 fractions, then concatenated into 12 by high-pH C18 reversed phase chromatography before analysis on an Orbitrap Fusion (Thermo Fisher Scientific) over the course of 2 h over 2–30% (v/v) acetonitrile in 0.1% (v/v) formic acid (Dionex UltiMate 3000) on a 250 × 0.075 mm column (Dr. Maisch Reprosil-PUR 1.9 μm). Spectra were matched against the IWGSC refseqV2 and the IBSC_V2 peptide databases by CometMS and results rescored and filtered to 2% peptide FDR in TPP V5.0. Details of spectra matching to the wheat and barley *THI4* genes are given in Supplementary Table 2.

### Bioinformatics

The NCBI dbEST database and the transcriptomes from the 1,000 Plants project (https://db.cngb.org/) [16] were searched using tblastn with the Arabidopsis THI4 protein as query, focusing specifically on presence/absence of an active-site Cys residue. Other resources used to search THI4 protein sequences were the NCBI non-redundant (nr) protein database (for ~500 plant sequences) and the THI4 subsystem of the SEED database [17] (for 114 microbial sequences). Sources of RNA-seq and THI4 sequence were as follows: barley cv. Morex: https://ics.hutton.ac.uk/morexGenes/index.html; wheat cv. Chinese Spring: https://wheat.pw.usda.gov/WheatExp/; oat (*Avena sativa*) genotype Ogle-C, developing seeds: [18]. Gene identifiers for barley and wheat are listed in Supplementary Table 3. Sequences were aligned with ClustalW or MultAlin and plotted with BoxShade (https://embnet.vital-it.ch/). Phylogenetic trees were built with Mega 5.2 [19] or Phylogeny.fr [20]. The Cys THI4 sequences included in the trees were those shown experimentally to have thiazole synthase activity [7,21-24].

### 3D structure modeling

Homology models of wheat, barley, and oat non-Cys THI4s were produced using SWISS-MODEL [25], inputting amino acid sequences corresponding to the DNA sequences in Supplementary Table 4 and the *M. igneus* non-Cys THI4 crystal structure (PDB 4Y4N) [3] as template (sequence identities 27-30%). The cereal THI4 homology models and yeast Cys THI4 (PBD 3FPZ) [1] were aligned to *M. igneus* THI4 using PyMOL (PyMOL Molecular Graphics System, Version 2.4.5 Schrödinger, LLC) to model the reaction intermediate glycine imine and iron atom in the active site clefts. Structural annotations and measurements were made using PyMOL. Active site residue conservation was determined by sequence alignments made with MultAlin.

### THI4 constructs for E. coli expression

Barley, wheat, and oat non-Cys THI4 coding sequences were recoded for expression in *E. coli* (Supplementary Table 4) and synthesized by GenScript (Piscataway, NJ). The Arabidopsis THI4 sequence was the native cDNA, obtained from the Arabidopsis Biological Resource Center (stock number U14240). Constructs were truncated at the positions shown in Supplementary Figure 1 to remove the plastid targeting peptide. Sequences with an added C-terminal hexahistidine tag were cloned between the XbaI and XhoI sites in pET28b using primers 11-18 (Supplementary Table 1).

### Expression of THI4s in E. coli

pET28b constructs were introduced into strain BL21(DE3) CodonPlus. This expression strain was used because pilot tests with wild type strain MG1655 showed no expression of the cereal THI4s. Cultures were grown at 37°C in LB medium until OD_600_ reached 0.8, then induced by adding isopropyl β-D-1-thiogalactopyranoside (IPTG, final concentration 1 mM), and incubated for another 14-16 h at 37°C. Cells were harvested by centrifugation (6 000****g****, 15 min, 4°C) and stored at −80°C. Proteins were extracted from cell pellets by sonicating in 100 mM potassium phosphate (pH 7.2) containing 2 mM β-mercaptoethanol. Protein was determined by the Bradford method. Protein samples (5 μg) were separated by SDS-PAGE on 15% gels; proteins were detected by Coomassie Blue staining or Western blotting using anti-His_6_-tag antibodies (Thermo Fisher Scientific MA1-21315).

### Functional complementation tests in E. coli

pET28b constructs were introduced into an *E. coli* BL21(DE3) Δ*thiG* strain constructed by recombineering [6,26]. Three independent clones of each construct were used to inoculate 3 ml of MOPS medium [27] containing 0.2% glucose, 100 nM thiamin, and 50 μg/ml kanamycin. After incubation at 37°C for 18 h, cells were harvested by centrifugation, washed five times with thiamin-free MOPS medium, resuspended in 500 μl of the same medium, serially diluted in ten-fold steps, and spotted on MOPS medium plates containing 0.2% glucose, 1 mM IPTG, plus or minus 1 mM Cys. Plates were then incubated at 37°C in aerobic conditions as described [6]. After 6 d (or 2 d for Arabidopsis), images were captured and digitized to quantify cell growth using ImageJ software. Briefly, an area of 155 × 155 pixels was chosen around the fourth serial dilution spot such that it contained the entire spot. ImageJ was then used to manually mark those parts of the area that contained colonies; the resulting values were then expressed as percent of the area occupied by colonies.

## RESULTS

### Arabidopsis THI4 exists mainly in its desulfurated form in planta

Because suicidal operation of Cys THI4s necessarily converts the active-site Cys residue to DHAla [1], the occurrence of DHAla at this position in a THI4 isolated from a plant would be diagnostic for a suicide mechanism. We therefore introduced a His_6_-tagged version of Arabidopsis *THI4* (synonyms: *THI1*, *TZ*), under the control of the native *THI4* promoter, into the Arabidopsis *tz-1* mutant. This mutant has a lethal thiazole synthesis defect attributable to the near-absence of *THI4* mRNA [11,23]. The *THI4* transgene complemented the *tz-1* mutation, allowing the transgenic plants to grow normally. Such homologous complementation of a *tz* mutant has not been obtained up to now [28,29]; it directly confirms the indirect evidence that the classical *TZ* locus encodes THI4 [23,28].

Proteins were isolated from the leaves of complemented plants and the His_6_-tagged THI4 protein was purified by Ni^2+^ affinity chromatography. The purified THI4 protein was then reduced and alkylated with acetamide, digested with trypsin, and analyzed by LC-MS/MS. We obtained peptide spectra with near-complete coverage of the protein sequence, including the two peptides predicted to contain Cys: 197-VGGVVTNWALVAQNHHTQSCMDPNVMEAK-225, which includes the active-site Cys216, and the downstream neighboring peptide 226-IVVSSCGHDGPFGATGVK-243 (Figure 2A). The ratio of spectral counts for these peptides and their corresponding desulfurated (i.e. DHAla-containing) versions showed that the shorter peptide was present mainly as the Cys (i.e. carbamidomethyl Cys) version (85% of spectra) while the longer peptide was present mainly as the desulfurated version (61% of spectra) (Figure 2B). To check that chemical modification of Cys was not biasing the ratio of spectra observed, trypsin digestion was also done in the absence of reduction and alkylation. Under these conditions the shorter peptide was again mainly observed in the Cys form (69% of spectra) while the longer peptide was almost exclusively in the desulfurated form (96% of spectra) (Figure 2B). The occurrence of a modest proportion of desulfurated Cys in the shorter peptide is attributable to desulfuration during sample workup [30]. Collectively, these data show that the majority of the THI4 protein extracted from Arabidopsis leaves had undergone sulfur loss from the active-site Cys, but not from a Cys residue elsewhere in the protein sequence. This difference between the two Cys residues is consistent with enzyme-mediated sulfur loss from the active-site Cys through the suicidal action of THI4, and with the absence of a repair pathway for desulfurated Arabidopsis THI4, as is the case for yeast THI4 [1].

**Figure 2.**
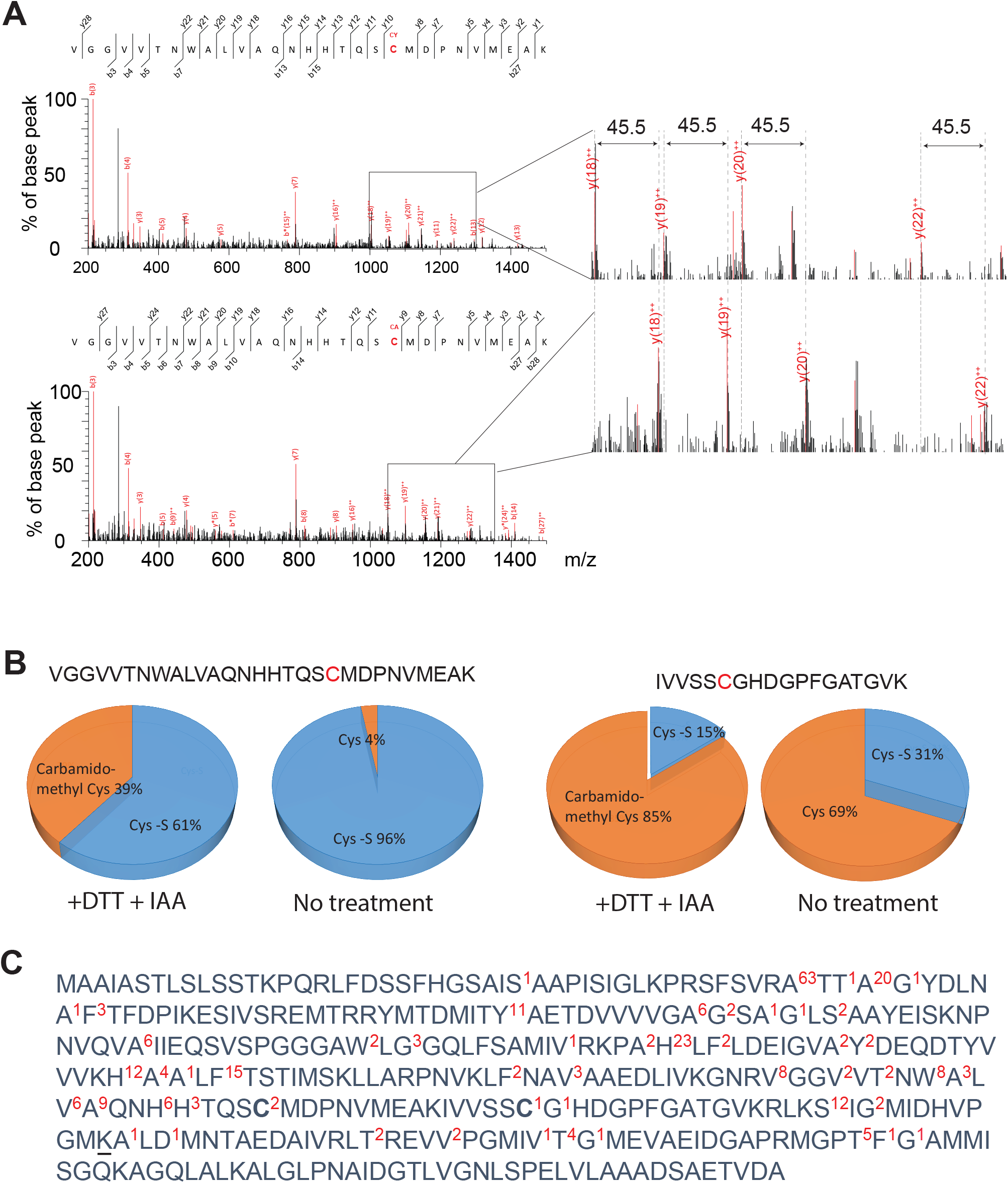
Characterization of Cys modifications in recombinant THI4 from Arabidopsis. (**A**) THI4 active-site Cys-containing peptide VGGVVTNWALVAQNHHTQSCMDPNVMEAK 4^+^-charged was characterized by mass spectrometry, with or without reduction with dithiothreitol (DTT) and alkylation with iodoacetamide (IAA). Two modifications occur at the Cys site: desulfuration (Cys —S) with 34 Da mass decrease and carbamidomethylation with a 57 Da mass increase. Pairs of double charged fragmentations (y-ion) show a 45.5 m/z shift between desulfurated and carbamidomethylated forms. (**B**) Spectral counting was used to quantify the relative proportion of desulfurated and carbamidomethylated (or free Cys) forms of the active-site Cys-containing peptide and a Cys-containing peptide from elsewhere in the protein. (**C**) A separate trypsin-digested THI4 protein sample without reduction and alkylation treatment was used to identify native N-termini and a GlyGly modification of Lys (K residue underlined) consistent with an ubiquitination site. Superscript numbers in red denote the number of spectra (with the criterion FDR <3%) determined by mass spectrometry.

Given the fast turnover of Arabidopsis THI4 [9], we ran semi-tryptic searches of the peptide spectra to identify novel N-termini that could indicate proteolytic sites. This showed a predominance of non-tryptic N-termini for the peptides 47-TTAGYDLNAFTFDPIK-62 and 50-GYDLNAFTFDPIK-62 (Figure 2C), consistent with the known N-terminal cleavage site of signal peptides by plastid processing enzymes after import of THI4 into the chloroplast [31]. However, this analysis also provided evidence of several cut sites in the middle of the THI4 protein, consistent with the action of Clp plastid proteases. We also found strong evidence for an ubiquitination site on the peptide 247-SIGMIDHVPGMKALDMNTAED-AIVR-271 based on a GlyGly modification of its Lys residue. This modification would be consistent with THI4 degradation via a combination of plastid proteolysis and subsequent targeting to the cytosolic proteasome.

### Bioinformatics indicates that cereals have grain-specific non-Cys THI4s

Searching the NCBI non-redundant protein database using Arabidopsis THI4 as query detected ~500 Cys THI4 sequences and two non-Cys THI4 sequences in plants. One non-Cys sequence (GenBank AAZ93636), from a cDNA from rice cv. Minghui 63 [24], is almost surely artifactual because the corresponding Minghui 63 genome sequence [32] encodes a canonical Cys THI4. The other non-Cys sequence (GenBank CDM85402), from wheat cv. Chinese Spring, is not artifactual as explained below.

To extend the search for non-Cys THI4 sequences we interrogated the NCBI dbEST expressed sequence tag (EST) database and the 1,000 Plants (1KP) transcriptome database [16] using tblastn with Arabidopsis THI4 as query. dbEST was found to contain wheat, barley, and oat THI4 sequences of two types: those encoding a canonical Cys THI4 similar to those of other flowering plants and those encoding a paralogous non-Cys THI4 in which His (wheat, barley) or Asp (oat) replaces the active-site Cys (Supplementary Figure 1). In all three species, the ESTs specifying non-Cys THI4s come only from developing grains while the Cys THI4 ESTs are from various organs (Supplementary Figure 2).

This EST evidence for non-Cys THI4s in cereals prompted searches of RNA-seq datasets, which corroborated the EST data and showed the timing of non-Cys THI4 expression during grain development. Thus, the non-Cys THI4 of wheat cv. Chinese Spring is expressed at the milk stage (Zadoks scale 75) (Figure 3A), which is ~10 d after pollination, near the mid-point of the grain filling period. Similarly, the non-Cys THI4 of barley cv. Morex is expressed at 15 d after pollination (Figure 3B), and a non-Cys THI4 is the main type of THI4 expressed in oat genotype Ogle-C from ~7 d to ~28 d after pollination (Supplementary Table 5) [18]. The expression of these non-Cys THI4s in the mid-to late stages of grain filling corresponds to when the central regions of the grain become hypoxic [33-35].

**Figure 3.**
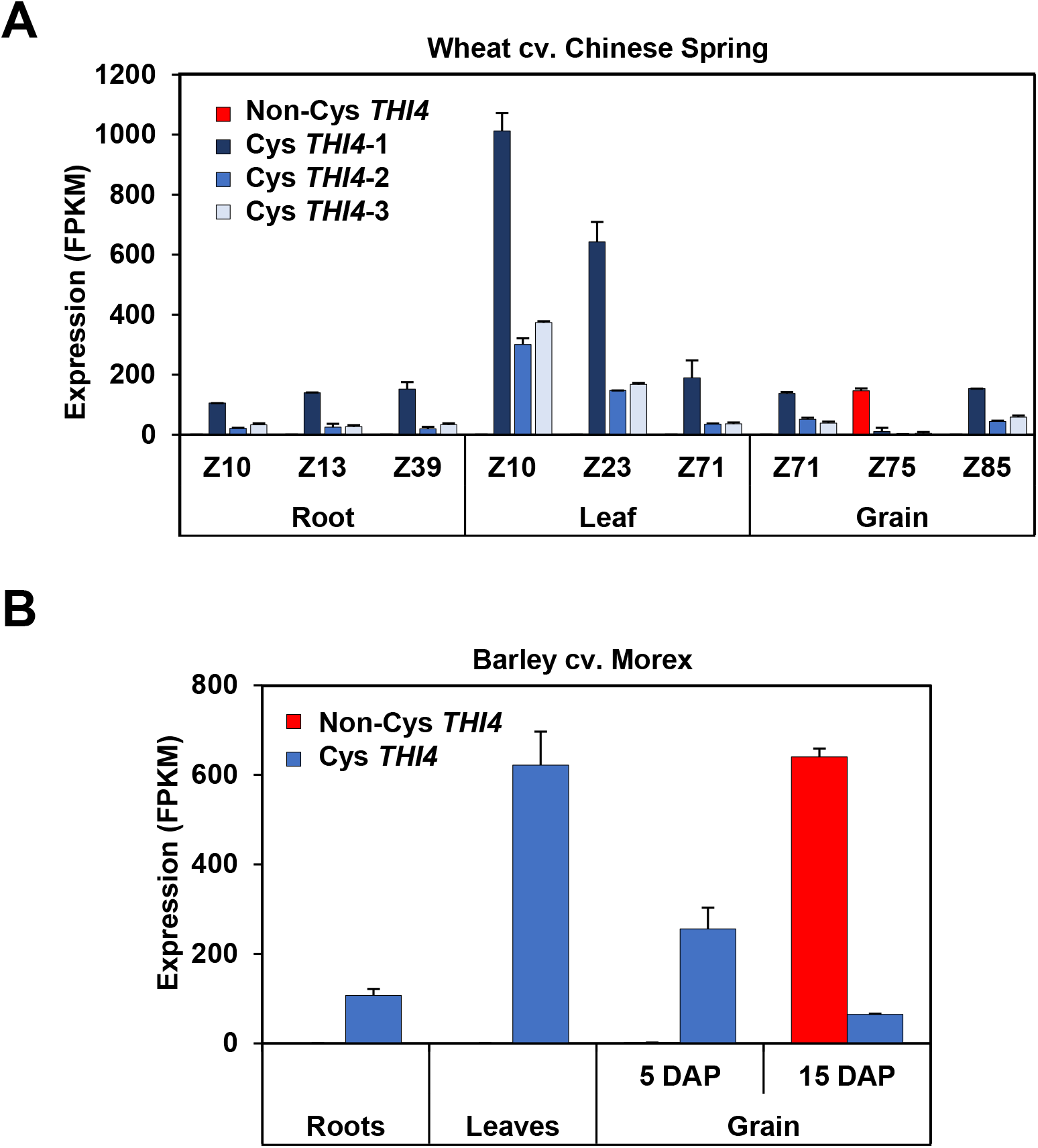
RNA-seq evidence for cereal non-Cys THI4s that are expressed in developing grains. (**A**) Expression of the single non-Cys and three Cys *THI4* genes of wheat cv. Chinese Spring in roots, leaves, and grains at various stages on the Zadoks (Z) growth stage scale. Values are means and s.e.m. of two replicates. Data are from https://wheat.pw.usda.gov/WheatExp/. (**B**) Expression of the single non-Cys and Cys THI4 genes of barley cv. Morex in roots, leaves, and grains at two developmental stages. DAP, days after pollination. Values are means and s.e.m. of three replicates. Data are from https://ics.hutton.ac.uk/morexGenes/index.html.

### Cereal non-Cys THI4 sequences diverge substantially from plant Cys THI4 sequences

Besides lacking an active-site Cys, cereal non-Cys THI4 sequences differ in other ways from the canonical Cys THI4 sequences of cereals and other plants (Supplementary Figure 1 and Supplementary Table 6). First, the non-Cys THI4s have substitutions at six positions that are fully conserved in plant Cys THI4s [6,7,36]. These substitutions are at Val^66^, Ala^91^, Gly^122^, Pro^236^, Gly^297^, and Met^303^ (numbered using the full length Arabidopsis THI4 sequence). Three of these residues (Gly^122^, Gly^297^, and Met^303^) are also fully conserved in prokaryote Cys and non-Cys THI4 sequences (Supplementary Figure 3). Other differences are that the cereal non-Cys THI4s have a one- or two-residue insertion after Pro^255^ and that wheat and oat non-Cys THI4s have small insertions after Thr^213^ (just upstream of the residue that replaces the active-site Cys). Lastly, non-Cys THI4s lack a ~20-residue C-terminal region that is conserved in monocot and eudicot Cys THI4s. Nothing is known about this region’s structure or function; the Arabidopsis THI4 crystal structure does not include it because the residues involved were missing from electron density maps [7] and the yeast and methanococcal THI4 proteins lack it [1,3,4].

The shared sequence features of wheat, barley, and oat non-Cys THI4s suggest a common evolutionary ancestor. Phylogenetic analyses support this possibility; independent sequence alignment and tree-drawing algorithms place the non-Cys THI4s in a distinct clade, separated from monocot and eudicot Cys THI4s (Supplementary Figure 4).

### Proteomics confirms that wheat and barley endosperms contain a non-Cys THI4

As the presence of endosperm mRNAs encoding a non-Cys THI4 does not prove that the protein is made, we sought validating proteomics evidence. Analysis of deep proteome profiling in wheat endosperm redundantly identified 16 THI4 tryptic peptides 318 times. These derive from four different genes, three of which have a Cys-containing active site and one a His-containing site (Figure 4A and Supplementary Table 2). Sixty-eight of these peptide spectra could be derived from active-site Cys-containing THI4 isoforms, seven of which could only be derived from the Cys-containing protein while the other 61 could be derived from either the Cys isoforms or the His-containing isoform. A total of 185 spectral matches were specific to one wheat isoform (TraesCS3B02G435500.1) predicted by RNA-seq and EST evidence (Figure 3A and Supplementary Figure 2A) to be present only in grains. This isoform has a His residue in the active site. The peptide 196-VTGVVTNWALVSMNQDTHSQTQSHMDANVM-EAK-228 containing this His active site sequence was experimentally observed in the wheat endosperm protein extracts. The large discrepancy in identification rates of peptides specific to the His-*vs.* Cys-containing isoforms (185 *vs*. seven) suggests that the His-containing THI4 predominates in wheat grains. A parallel analysis conducted for developing barley grains (Figure 4B and Supplementary Table 2) showed a similar situation with 189 spectra deriving specifically from the His-containing THI4 and just five from the Cys-containing THI4. As for wheat, the characteristic His-containing active-site peptide was experimentally observed. The huge (>20-fold) difference in apparent abundance between the Cys- and His-containing THI4s, along with the transcriptome data (Figure 3 and Supplementary Figure 2), implies that the His-containing THI4 functionally replaces the canonical Cys THI4 in wheat and barley endosperm and therefore that the His-containing THI4 would need to provide any thiazole synthase activity required in this tissue. This inference, however, rests on the assumption that wheat and barley endosperm express a complete thiamin synthesis pathway because, in the absence of the rest of the pathway, there would be no obvious need for a thiazole synthase [37].

**Figure 4.**
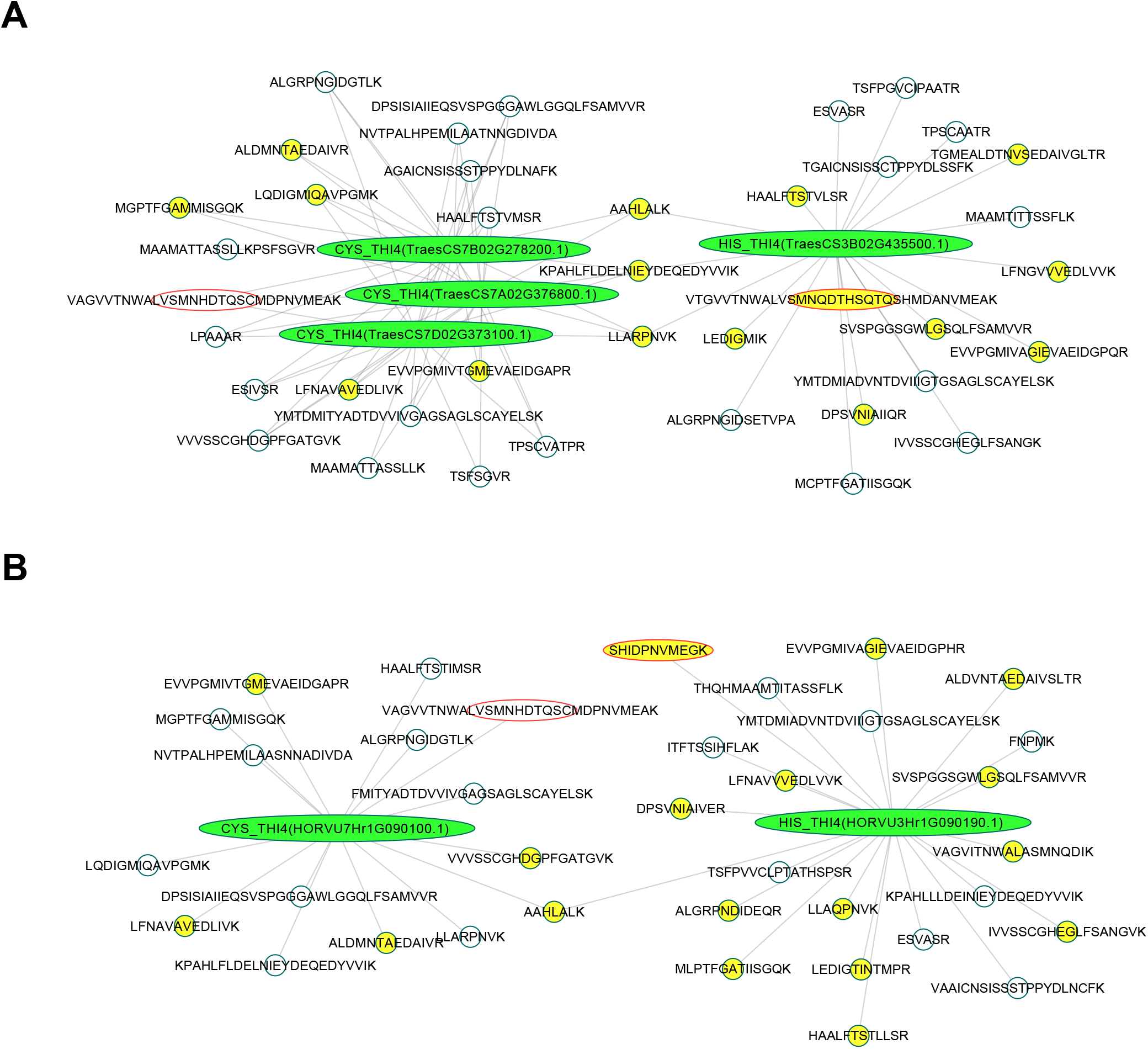
Tryptic peptides for THI4 in the wheat and barley genomes and peptide identifications in seeds. (**A**) Peptides identified in proteomic profiling of the endosperm from mature wheat grains (cv. Wyalkatchem). Four genes (green ellipses) were identified in the wheat genome by searching with the Arabidopsis THI4 (At5g54770) protein sequence. All possible tryptic peptides longer than five residues which could be derived from these proteins are shown (grey circles) with those positively identified in wheat seeds shown in yellow. Spectra matching to peptides from the Cys-containing isoform were identified 68 times and the His-containing isoform 246 times. Details of these spectra are given in Supplementary Table 2. (**B**) Peptides identified in proteomic profiling of developing endosperm (late dough stage) from barley (cv. Morex). Two genes (green ellipses) were identified in the barley genome by searching with the Arabidopsis THI4 protein sequence. Spectra matching to the Cys-containing THI4 were identified seven times while spectra matching to the His-containing THI4 were identified 191 times. Details of these spectra are given in Supplementary Table 2.

### Transcriptome data indicate that cereal grains have a complete thiamin synthesis pathway

To validate the assumption that wheat, barley, and oat grains express all the genes (besides *THI4*) needed to synthesize thiamin *de novo*, as is the case for maize endosperm [38], we searched wheat and barley RNA-seq datasets for expression data on *THIC* (which makes the pyrimidine moiety of thiamin) and *TH1* (the bifunctional enzyme that phosphorylates the pyrimidine moiety and couples it to the thiazole moiety). Both types of genes are expressed in developing grains, including endosperm tissue, at levels broadly comparable to those in leaves (Supplementary Figures 5 and 6). As leaves are a major site of thiamin synthesis [38,39], this transcriptomic evidence supports the inference that thiamin is synthesized in cereal grains using a non-Cys THI4 as the thiazole synthase.

### 3D Structural models are consistent with non-Cys THI4s being catalytic thiazole synthases

Nine of the active site residues annotated in the yeast, Arabidopsis, and *M. igneus* THI4 crystal structures (PDB 1RP0, 3FPZ, and 4Y4N, respectively) are identical or similar to each other (Supplementary Figure 3). Of these nine residues, seven – including the Fe^2+^-binding Asp and Glu residues – are conserved or conservatively replaced in all three cereal non-Cys THI4 sequences (Supplementary Figure 3). This high level of conservation enabled construction of 3D structural models of the active sites of the cereal THI4s with the glycine imine reaction intermediate [3] bound (Figure 5 and Supplementary Figure 7). The models indicated that the cereal THI4 active sites are similar in size and overall configuration to those of both the *M. igneus* non-Cys THI4 (Figure 5A) and the yeast Cys THI4 (Figure 5B), and that the conserved Fe^2+^-binding Asp and Glu residues in the cereal THI4s are appropriately positioned to mediate the sulfur transfer reaction (Figure 5C,D,E). The models also indicated that neither of the Cys residues in the active site cleft of the cereal THI4s is a plausible sulfur source because they are ≥13 Å distant from the carbon atom to which the sulfur is attached (Supplementary Figure 7). Furthermore, one of these Cys residues is also present in Arabidopsis THI4 (Figure 2 and Supplementary Figure 3). Thus, modeling revealed several structural features of cereal non-Cys THI4s that are consistent with catalytic thiazole synthase activity, and none that are inconsistent with it.

**Figure 5.**
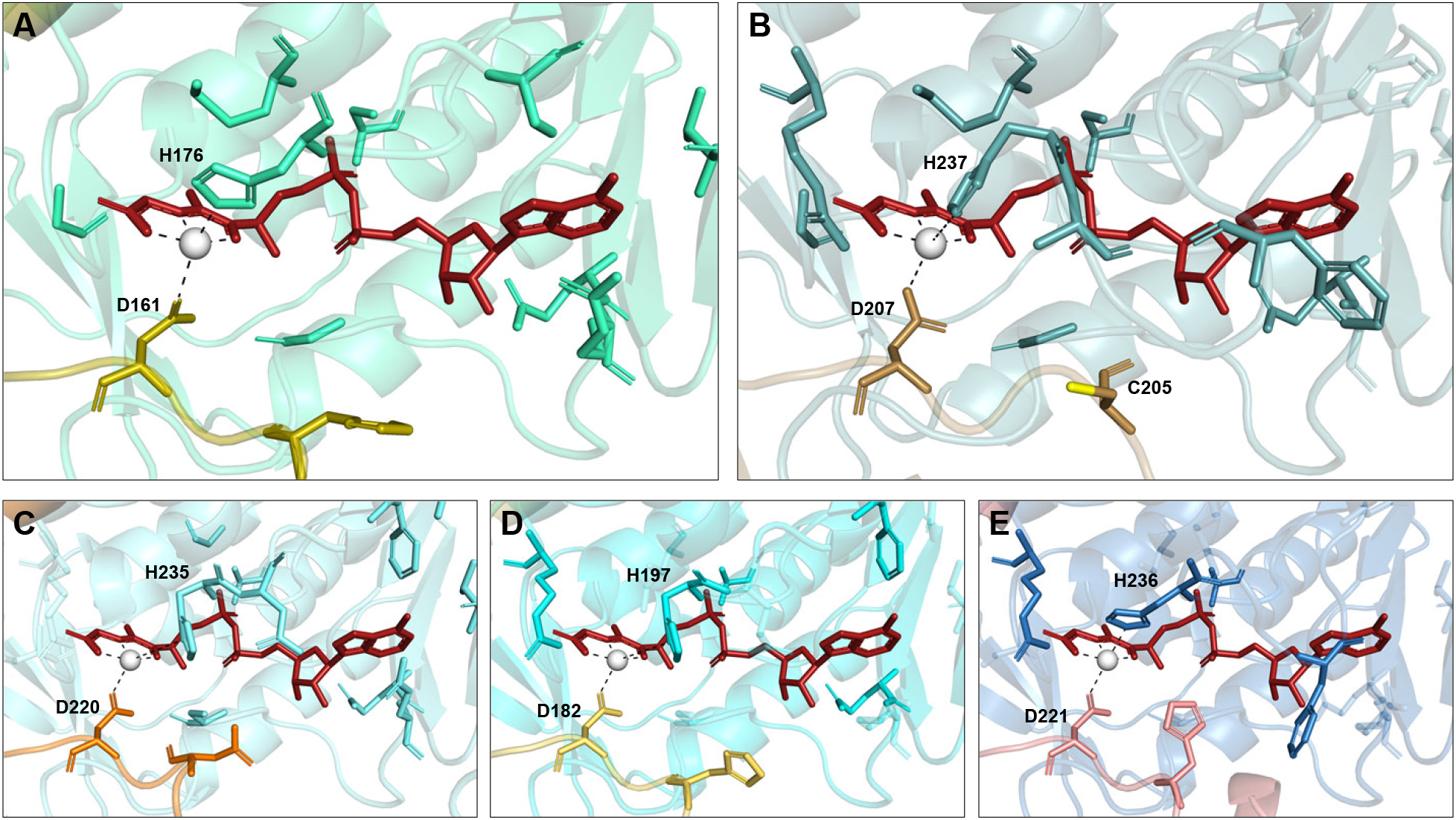
Modeling of the active site clefts of cereal non-Cys THI4s. All structures include bound glycine imine reaction intermediate (red) and Fe^2+^ (gray sphere); the two Fe^2+^-binding residues are indicated. (**A**) *M. igneus* non-Cys THI4 (PDB 4Y4N) co-crystallized with glycine imine (brick-red) and Fe^2+^ (gray sphere). (**B**) Yeast THI4 (PDB 3FPZ) with the sulfur atom of the sulfur-donating Cys (C205) in yellow. (**C**) Oat, (**D**) barley, and (**E**) wheat non-Cys THI4 homology models (template: 4Y4N, SWISS-MODEL) with glycine imine and Fe^2+^ modeled into the active sites using Pymol 2.3.5 align command. Putative active site residues predicted by MultAlin and Pymol sequence alignments are shown in stick format (non-identical residues are shown at 0.7 transparency). Putative residue-Fe^2+^ interact-ions predicted to be within 3 Å are indicated by black dashes. In cereal non-Cys THI4s, the loop that contains C205 in yeast THI4 more closely resembles the corresponding loop in *M. igneus* non-Cys THI4.

### Functional complementation confirms that non-Cys THI4s have thiazole synthase activity

Given the above indirect evidence that non-Cys THI4s can functionally replace Cys THI4s, we tested His_6_-tagged non-Cys THI4s for ability to complement the thiazole auxotropy of a Δ*thiG* mutant of *E. coli* strain BL21(DE3). This engineered expression strain was used because wild type strain MG1655 showed no detectable expression of non-Cys THI4s in pilot studies. All three THI4s showed expression in strain BL21(DE3) that was detectable by Western blot, the band in wheat being stronger than in barley or oat; wheat also gave a strong Coomassie-stained protein band (Supplementary Figure 8A). Complementation tests were run in aerobic conditions, without or with Cys supplementation to raise the intracellular sulfide level [40]. Tests could not be run in anaerobic conditions because strain BL21(DE3) has major defects in anaerobic metabolism [41]. When Cys was present, all three non-Cys THI4s showed significant complementing activity, albeit far lower that of the Arabidopsis THI4 included as a benchmark (Figure 6 and Supplementary Figure 8B). The difference from Arabidopsis was not due to toxicity of the cereal THI4s because, when given thiamin, strains expressing cereal THI4s grew as well as those expressing Arabidopsis THI4 (Supplementary Figure 8B). None of the cereal THI4s had complementing activity when Cys was absent (Figure 6). The dependence of complementation on Cys supplementation is consistent with sulfide being the sulfur donor in the cereal THI4 reaction.

**Figure 6.**
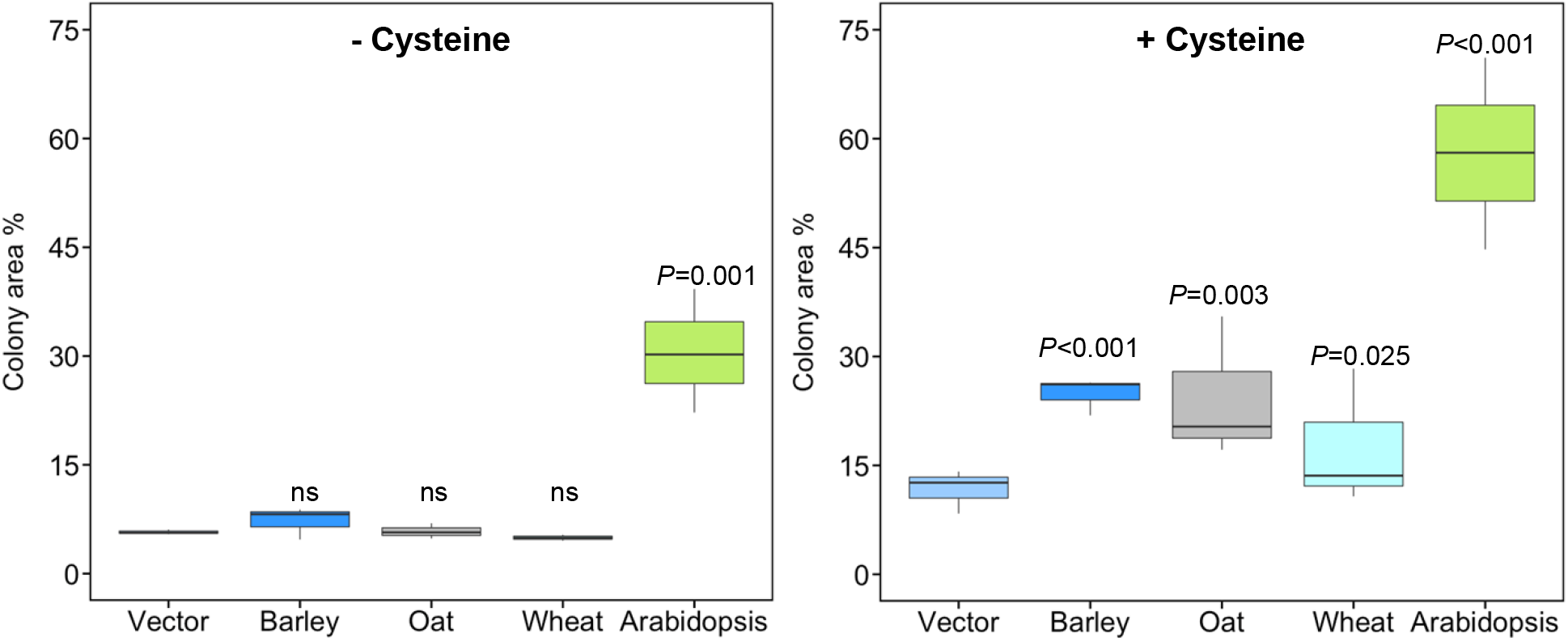
Complementation of an E. coli thiazole auxotroph by cereal non-Cys THI4s. Cells of an *E. coli* BL21(DE3) Δ*thiG* strain harboring the pET28 vector alone (Vector) or containing a His_6_-tagged cereal no-Cys *THI4* gene (or the Arabidopsis Cys *THI4* gene as a benchmark) were cultured in aerobic conditions in MOPS medium containing 0.2% glucose, 1 mm IPTG, minus or plus 1 mM Cys. Overnight liquid cultures of three independent clones of each construct were serially diluted in ten-fold steps and spotted on plates. After incubation at 37°C for 6 d (or 2 d for Arabidopsis), images were captured and the images of the fourth dilution in the series were digitized to quantify growth (as percent of the dilution spot occupied by colonies). The images are shown in Supplementary Figure 8B. Statistical significance was determined on log_10_-transformed data by Student’s *t*-test. ns, non-significant.

## DISCUSSION

### The evidence for both suicidal and catalytic plant THI4s

We report here direct proteomic evidence that the canonical Cys THI4s of plants are suicide enzymes. We also present *prima facie* evidence that certain cereal THI4s cannot obtain the sulfur atom needed for thiazole synthesis from an active-site Cys residue because they lack such a residue. These non-Cys THI4s are thus potentially truly catalytic thiazole synthases that mediate multiple turnovers.

The evidence that cereal non-Cys THI4s are functional catalytic thiazole synthases is four-fold: (i) non-Cys catalytic thiazole synthases are well precedented among prokaryotes; (ii) cereal non-Cys THI4s largely replace canonical Cys THI4s in the thiamin biosynthesis pathway in grains; (iii) the amino acid sequences and predicted structures of cereal non-Cys THI4s point to catalytic thiazole synthase activity; and (iv) non-Cys THI4s have detectable ability to complement thiazole auxotrophy in *E. coli* and this complementation is dependent on an exogenous sulfur source. While coherent, this evidence is challengeable. One challenge concerns the complementation data: cereal non-Cys THI4s all had far lower complementing activity than Arabidopsis THI4 (Figure 6 and Supplementary Figure 8B) and wheat non-Cys THI4 was no more active than those of barley or oat despite its higher expression level (Supplementary Figure 8A). To which we respond (i) that poor availability or insertion of the metal cofactor [3], a bad truncation point (Supplementary Figure 1), the added His_6_-tag, and incorrect folding [42] could all lower the activity of cereal THI4s in *E. coli*, and (ii) that solubility does not necessarily predict activity because recombinant proteins often form soluble but catalytically inactive aggregates in *E. coli* [43,44]. Although we set out to confirm the complementation results by *in vitro* THI4 activity assays [1,4], the weak recombinant expression of barley and oat THI4s and the probable inactivity of the expressed wheat THI4 protein (Supplementary Figure 8A) made this approach impracticable.

Another challenge is that the cereal non-Cys THI4s might not be thiazole synthases at all, given the proposed moonlighting functions of other plant THI4s in mitochondrial DNA damage tolerance [45] and in anion transport across the plasma membrane of guard cells [46]. In this view, the primary activity of cereal non-Cys THI4s would be the moonlighting one and the complementing activity would be secondary and residual. To which we respond that the proposed moonlighting roles are most probably artifacts because (i) the evidence supporting them comes from unphysiological systems, notably expression in a heterologous host (*E. coli*) [45] or overexpression of a THI4-green fluorescent protein fusion *in planta* [46], and (ii) roles in mitochondria or the plasma membrane are controverted by strong evidence that *native* THI4 is localized only in plastids [29,47].

If cereal non-Cys THI4s are indeed thiazole synthases, their occurrence and their localization in developing grains raise questions about their evolutionary origin, their adaptive significance, and their possible relationship to oxygen. We discuss these points in sequence below.

### Cereal non-CysTHI4s are probably of ancient origin and may be present in other grasses

That the cereal non-Cys THI4s fall into the same clade (Supplementary Figure 4) implies a single origin. As the grass tribes to which wheat and barley (Triticeae) and oat (Poeae) belong are estimated to have diverged ~30 Myr ago [48], this origin was likely ancient and long preceded domestication. In relation to the origin, it is worth noting that a mutant plant THI4 with an active-site Ala in place of Cys had thiazole synthase activity in anaerobic, high-sulfide conditions [6]. This finding suggests that, in an ancestral THI4 expressed during grain development, a mutation that replaced the active-site Cys with another residue could have launched the non-Cys THI4 clade, none of the other mutations in non-Cys THI4s (Supplementary Figure 1 and Supplementary Table 6) being a necessary precondition. The birth of the non-Cys THI4 clade might thus have been an easy one.

If non-Cys THI4s originated prior to the split between the Poeae (the largest grass tribe) and the Triticinae (also a fairly large tribe), they could well be present in many other grasses. And indeed, searching the draft genome sequence of rye (*Secale cereale*) [49] uncovered a non-Cys THI4 (Lo7_v2_scaffold_171684 3R) as well as a Cys THI4 (Lo7_v2_scaffold_527772 7R). The more grass genomes and transcriptomes come online, the more non-Cys THI4s will probably emerge.

### Catalytic THI4s could contribute substantially to grain ATP budgets

Given the likelihood that cereal non-Cys THI4s are catalytic, we estimated the energetic benefit to a developing grain of replacing a suicide THI4 with a catalytic one. Analogous estimates for vegetative tissues show a substantial benefit of this replacement [10]. We considered the central region of the developing grain in mid-grain fill because this region is hypoxic and has a low ATP level that can limit starch synthesis [34,50]. Because starch synthesis from imported sucrose is the predominant metabolic flux in the endosperm [51,52] we used the ATP cost of this flux as a benchmark. Taking sucrose to be metabolized via sucrose synthase, we obtained a value of 4.3 μmol ATP d^−1^ per grain (Table 1). The estimated cost of turnover of a Cys THI4 was 0.75 μmol ATP d^−1^ per grain, i.e. 17% of the starch synthesis cost (Table 1). This estimate assumes a half-life of thiamin in endosperm equal to that in leaves, but even were the half-life of thiamin in endosperm several-fold longer, THI4 turnover would still represent a sizeable drain on the ATP budget. A catalytic THI4 could eliminate nearly all this drain, supposing that, like typical enzymes [53], it performs many reaction cycles before being degraded.

**Table 1.** Comparison of the estimated ATP costs of turnover of a suicidal Cys THI4 and of starch synthesis from sucrose for cereal endosperm in mid-grain fill

| Parameter (units) | Values | References |
| --- | --- | --- |
| Grain dry weight (DW) (mg) | 20 | [56] |
| Total thiamin content<sup>a</sup> (nmol per grain) | 0.2 | [57] |
| Thiamin turnover rate (d<sup>-1</sup>) | 1.7 | [10] |
| Thiamin and THI4 synthesis rates (nmol d<sup>-1</sup> per grain) | 0.34 |  |
| ATP cost of THI4 turnover<sup>b</sup> (μmol d<sup>-1</sup> per grain) | 0.75 | [10,58] |
| Starch synthesis rate<sup>c</sup> (mg d<sup>-1</sup> per grain) | 1.4 | [34,56] |
| Rate of hexose unit addition to starch (μmol d<sup>-1</sup> per grain) | 8.6 |  |
| ATP cost of hexose unit addition<sup>d</sup> (μmol d<sup>-1</sup> per grain) | 4.3 | [51,52] |
| Cost of THI4 turnover (% of starch synthesis) | **17** |  |

<sup>a</sup> Approximate average value for wheat and barley grains.
<sup>b</sup> For a 351-residue barley Cys THI4 @ turnover cost of 6.3 ATP per residue.
<sup>c</sup> Assuming grain starch content = 60% of dry weight.
<sup>d</sup> Assuming synthesis of starch from sucrose via sucrose synthase and full reclamation of the phosphate bond energy in the pyrophosphate formed by ADP-glucose pyrophosphorylase.

The above estimate makes energy-saving a plausible driver for evolution of a catalytic, sulfide-utilizing THI4 in the oxygen-deficient, ATP-limited endosperm of barley, wheat, and oat. It also warrants equipping rice and maize with such enzymes as an experimental yield-enhancement strategy.

### An oxygen connection?

There is an intriguing parallel between the observations that prokaryotic non-Cys THI4s are largely confined to anaerobes and microaerophiles [4,6] and that cereal non-Cys THI4s are expressed mainly in developing grains at stages when hypoxia occurs [50]. Could oxygen sensitivity of the thiazole synthase reaction explain the parallel? In both non-Cys and Cys THI4s, an Fe^2+^ ion participates in the sulfur transfer reaction [3,4]. However, Fe^2+^ is oxygen-sensitive, being rapidly oxidized to Fe^3+^ and then precipitating [54]. Additionally, non-Cys THI4s use HS^−^, which is also oxygen-sensitive and readily undergoes oxidation [55]. The Cys THI4s of yeast, plants, and the aerobic archaeon *Haloferax volcanii* have clearly mastered the chemical challenge of running the suicidal thiazole synthase reaction efficiently in aerobic conditions *in vivo* [3,36], although how they cope with oxygen is not clear [1,3,22]. Conversely, most prokaryotic non-Cys THI4s seem not to have mastered, or fully mastered, the challenge of running the HS^−^-dependent catalytic reaction in aerobic conditions [4,6], perhaps due to the extra oxygen sensitivity entailed by using HS^−^ as sulfur donor. If cereal non-Cys THI4s have likewise not learned the trick of working well in air, it would explain why they are expressed only in hypoxic developing grains – and possibly also why they have low complementing activity in aerobic conditions.

## Acknowledgments

We thank M.J. Ziemak for technical support.

## Author Contribution

A.D.H., J.J., G.A.W.B., and A.H.M. designed the research; J.J., G.A.W.B., J.A.P., and J.-D.G.-G. performed microbiological and biochemical experiments with supervision from A.D.H.; C.B. and L.-Y.C. carried out bioinformatics analyses with supervision from J.A.P. and A.D.H; L.L., O.D., and A.H.M. carried out proteomics analyses; J.J. and A.D.H. wrote the article.

