## Supplementary Material for "Bioinformatic and experimental evidence for suicidal and catalytic plant THI4s"

```
                                                                           *
Arabidopsis      1       MAAIASTLSLSSTKPQRLFDSSFHGSAISAAPI--SIGLKPRSFSVRATTAGYDLNAFTFDPIKESIVSREMTR
Alder Cys        1   MASTLTSKPQRLALFENSASSFYGTPLAPSSIRVQPT--KAGAKPSISMSGAPSPPYDLKAFTFDPIKESIVSREMTR
Barley Cys       1 MAAMATTASSLLKSSFSGVRLPAAARTPS----CVATPR--AG-AICNSIS--SSTPPYDLNAFKFSPIKESIVSREMTR
Wheat Cys        1    MATTASSLLKTSFSGVRLPAAARTPS----CVATPR--AG-AICNSIS--SSTPPYDLNAFKFSPIKESIVSREMTR
Oat Cys          1 MAAMATTASSLLKTSFVGAHLPAASRTPS----CMAVPR--AG-AICNSIS--SSTPPYDLNAFNFSPIKESIVSREMTR
Rice Cys         1 MAAMATTASSLLKTSFAGARLPAAARNPT----VSVAPR--TGGAICNSISSSSTPPYDLNAIRFSPIKESIVSREMTR
Maize THI1-1     1   MATAAASSLLKSSFAGSRLPAATRTTPASLVVATGPRGAGAGPICASMSMSSSNPPYDLTSFRFSPIKESIVSREMTR
Maize THI1-2     1   MATTAASSLLKSSFAGSRLPSATRTTTPSSVAVATPR-AGGGPIRASIS-SSPTPPYDLTSFRFSPIKESIVSREMTR
Barley non-Cys   1 MAAMTITASSFLKTSFPVVCLPTATHSPSRASTRRV-------AAICNSIS--SSTPPYDLNCFKFNPMKESVASREMTR
Wheat non-Cys    1 MAAMTITTSSFLKTSFPGVCIPAATRTPSCAATRRT-------GAICNSIS--SCTPPYDLSSFKFSPMKESVASREMIR
Oat non-Cys      1 MAAMATAAPILLKTSFAGACLPKAARSLTLDAVCTP-----RAGAIYNSIS--SSTPPYDFNAVKFKPIKESIASREMIR

Arabidopsis     73 RYMTDMITYAETDVVVVGAGSAGLSAAYEISKNPNVQVAIIEQSVSPGGGAWLGGQLFSAMIVRKPAHLFLDEIGVAYDE
Alder Cys       77 RYMMDMITYADTDVVVVGAGSSGL-VCYELSKNPSVQVAIIEQSVSPGGGAWLGGQLFSGMVVRKPAHLFLDELGIEYDE
Barley Cys      72 RYMTDMITYADTDVVIVGAGSAGLSCAYELSKDPSISIAIIEQSVSPGGGAWLGGQLFSAMVVRKPAHLFLDELNIEYDE
Wheat Cys       69 RYMTDMITYADTDVVIVGAGSAGLSCAYELSKDPSISIAIIEQSVSPGGGRWLGAQLFSAMVVRKPAHLFLDELNIEYDE
Oat Cys         72 RYMTDMITYADTDVVIVGAGSAGLSCAYELSKDPSISIAIIEQSVSPGGGAWLGGQLFSAMVVRKPAHLFLDELNIEYDE
Rice Cys        75 RYMTDMITYADTDVVVVGAGSAGLSCAYELSKDPSVSVAVIEQSVSPGGGAWLGGQLFSAMVVRKPAHLFLDELGVAYDE
Maize THI1-1    79 RYMTDMITYADTDVVIVGAGSAGLSCAYELSKDPAVSIAIVEQSVSPGGGAWLGGQLFSAMVVRKPAHLFLDELGVAYDE
Maize THI1-2    77 RYMTDMITHADTDVVIVGAGSAGLSCAYELSKDPTVSVAIVEQSVSPGGGAWLGGQLFSAMVVRRPAHLFLDELGVGYDE
Barley non-Cys  72 RYMTDMIADVNTDVIIIGTGSAGLSCAYELSKDPSVNIAIVERSVSPGGSGWLGSQLFSAMVVRKPAHLLLDEINIEYDE
Wheat non-Cys   72 RYMTDMIADVNTDVIIIGTGSAGLSCAYELSKDPSVNIAIIQRSVSPGGSGWLGSQLFSAMVVRKPAHLFLDELNIEYDE
Oat non-Cys     74 RYMADMITFSDTDVVIIGTGPAGLSCAYELSKDPSINIAIIEQSVSPGGSAWLGGQFCSAMVVRKTAHLFLDELNIPYDE

                                  †                      ‡
Arabidopsis    153 QDTYVVVKHAALFTSTIMSKLLARPNVKLFNAVAAEDLIVKGNRVGGVVTNWALVAQNHHT----QSCMDPNVMEAKIVV
Alder Cys      156 QDNYVVIKHAALFTSTIMSKLLARPNVKLFNAVAAEDLIVKGGRVGGVVTNWALVSMNHDT----QSCMDPNVMEAKVVV
Barley Cys     152 QEDYVVIKHAALFTSTIMSRLLARPNVKLFNAVAVEDLIVKENRVAGVVTNWALVSMNHDT----QSCMDPNVMEAKVVV
Wheat Cys      149 QEDYVVIKHAALFTSTVMSRLLARPNVKLFNAVAVEDLIVKEDRVAGVVTNWALVSMNHDT----QSCMDPNVMEAKVVV
Oat Cys        152 QEDYVVIKHAALFTSTVMSRLLARPNVKLFNAVAVEDLIVKENRVAGVVTNWALVSMNHDT----QSCMDPNVMEAKVVV
Rice Cys       155 QEDYVVIKHAALFTSTVMSRLLARPNVKLFNAVAVEDLIVKEGRVGGVVTNWALVSMNHDT----QSCMDPNVMESRVVV
Maize THI1-1   159 AEDYVVIKHAALFTSTVMSLLLARPNVKLFNAVAVEDLIVRGGRVGGVVTNWALVSMNHDT----QSCMDPNVMEAKVVV
Maize THI1-2   157 AEDYVVVKHAALFTSTVMSRLLARPNVKLFNAVAVEDLIVRRGRVGGVVTNWALVSMNHDT----QSCMDPNVMEAKVVV
Barley non-Cys 152 QEDYVVIKHAALFTSTLLSRLLAQPNVKLFNAVVVEDLVVKENRVAGVITNWALASMNQDI----KSHIDPNVMEGKIVV
Wheat non-Cys  152 QEDYVVIKHAALFTSTVLSRLLARPNVKLFNGVVVEDLVVKEHRVTGVVTNWALVSMNQDTHSQTQSHMDANVMEAKIVV
Oat non-Cys    154 QEDYVVIKHAALFTSTILSRLLTCPNVKLFNAVEVEDLIIKENHVAGVVTNWALVSMKHDTQ---SYDMDPNIMEAKVVV

Arabidopsis    229 SSCGHDGPFGATG--VKRLKSIGMIDHVP--GMKALDMNTAEDAIVRLTREVVPGMIVTGMEVAEIDGAPRMGPTFGAMM
Alder Cys      232 SSCGHDGPFGATG--VKSLRSIGMIDTVP--GMKALDMNVAEDAIVRLTREIVPGMIVTGMEVAEIDGAPRMGPTFGAMM
Barley Cys     228 SSCGHDGPFGATG--VKRLQDIGMIQAVP--GMKALDMNTAEDAIVRLTREVVPGMIVTGMEVAEIDGAPRMGPTFGAMM
Wheat Cys      225 SSCGHDGPFGATG--VKRLQDIGMIQAVP--GMKALDMNTAEDAIVRLTREVVPGMIVTGMEVAEIDGAPRMGPTFGAMM
Oat Cys        228 SSCGHDGPFGATG--VKRLQDIGMIETVP--GMKALDMNTAEDAIVRLTREVVPGMIVTGMEVAEIDGAPRMGPTFGAMM
Rice Cys       231 SSCGHDGPFGATG--VKRLQDIGMIDAVP--GMRALDMNTAEDEIVRLTREVVPGMIVTGMEVAEIDGAPRMGPTFGAMM
Maize THI1-1   235 SSCGHDGPFGATG--VKRLQDIGMISAVP--GMKALDMNTAEDEIVRLTREVVPGMIVTGMEVAEIDGAPRMGPTFGAMM
Maize THI1-2   233 SSCGHDGPFGATG--VKRLQDIGMISAVP--GMKALDMNAAEDEIVRLTREVVPGMIVTGMEVAEIDGAPRMGPTFGAMM
Barley non-Cys 228 SSCGHEGLFSANG--VKRLEDIGTINTMPR--MKALDVNTAEDAIVSLTREVVPGMIVAGIEVAEIDGPHRMLPTFGATI
Wheat non-Cys  232 SSCGHEGLFSANGKGVKRLEDIGMIKTVPRTGMEALDTNVSEDAIVGLTREVVPGMIVAGIEVAEIDGPQRMCPTFGATI
Oat non-Cys    231 SACGHDRKFGATI--VKHFQD-RMIETMPP-GMSVLDENMSEEDIVHYTREVVPGLIVTGIQVADIEGAPRMVPTFGATM

Arabidopsis    305 ISGQKAGQLALKALGLPNAIDGTLVGNL----SPELVLAADSAETVDA
Alder Cys      308 ISGQKAAHLALKALGLPNALDGSYVGGI----HPELILAADSAEIADA
Barley Cys     304 ISGQKAAHLALKALGRPNGIDGTLKNVTPAL-HPEMILAASNNADIVDA
Wheat Cys      301 ISGQKAAHLALKALGRPNGIDGTLKNVTPAL-HPEMILAATNNGDIVDA
Oat Cys        304 ISGQKAAHLALKALGRPNGIDGTLKNVTPAL-HPEMVLASADNGDIVDA
Rice Cys       307 ISGQKAAHLALKALGRPNAIDGTIKKAAAAAAHPELILASKDDGEIVDA
Maize THI1-1   311 ISGQKAAHLALKALGRPNAVDGTM---SPPL-REELMIAYKDD-EVVDA
Maize THI1-2   309 ISGQKAAHLALKALGRPNAVDGTIPEVSPAL-REEFVIASKDD-EVVDA
Barley_non-Cys 304 ISGQKAAHLALKALGRPNDIDEQRYSRESRRYTRRLS
Wheat non-Cys  312 ISGQKAAHLALKALGRPNGIDSETVPA
Oat non-Cys    307 ISGQKAAHLALKALGRPNSIDRTKRCTRS
```

**Supplementary Figure 1. Alignment of Cys and non-Cys THI4 sequences from barley, wheat, and oat with functionally validated Cys THI4 sequences from other angiosperms.**

The active-site Cys residue in each sequence shaded blue; residues that replace it are shaded red. Residues changed to the start Met in truncated versions of THI4 proteins are shaded green. The asterisk marks the experimentally determined N-terminus of Arabidopsis THI4 [31]. Substitutions at residues that are fully conserved in plant Cys THI4s are shaded in magenta. †, Ala residue whose replacement by Val in Arabidopsis THI4 abolished activity [28]. ‡, Val residue whose replacement by Met in maize THI1-2 abolished activity [22]. For the Arabidopsis THI4 protein, residue numbers in the crystal structure [PDB: 1RP0] are obtained by subtracting 44 from the numbers for the full-length protein shown in the alignment.

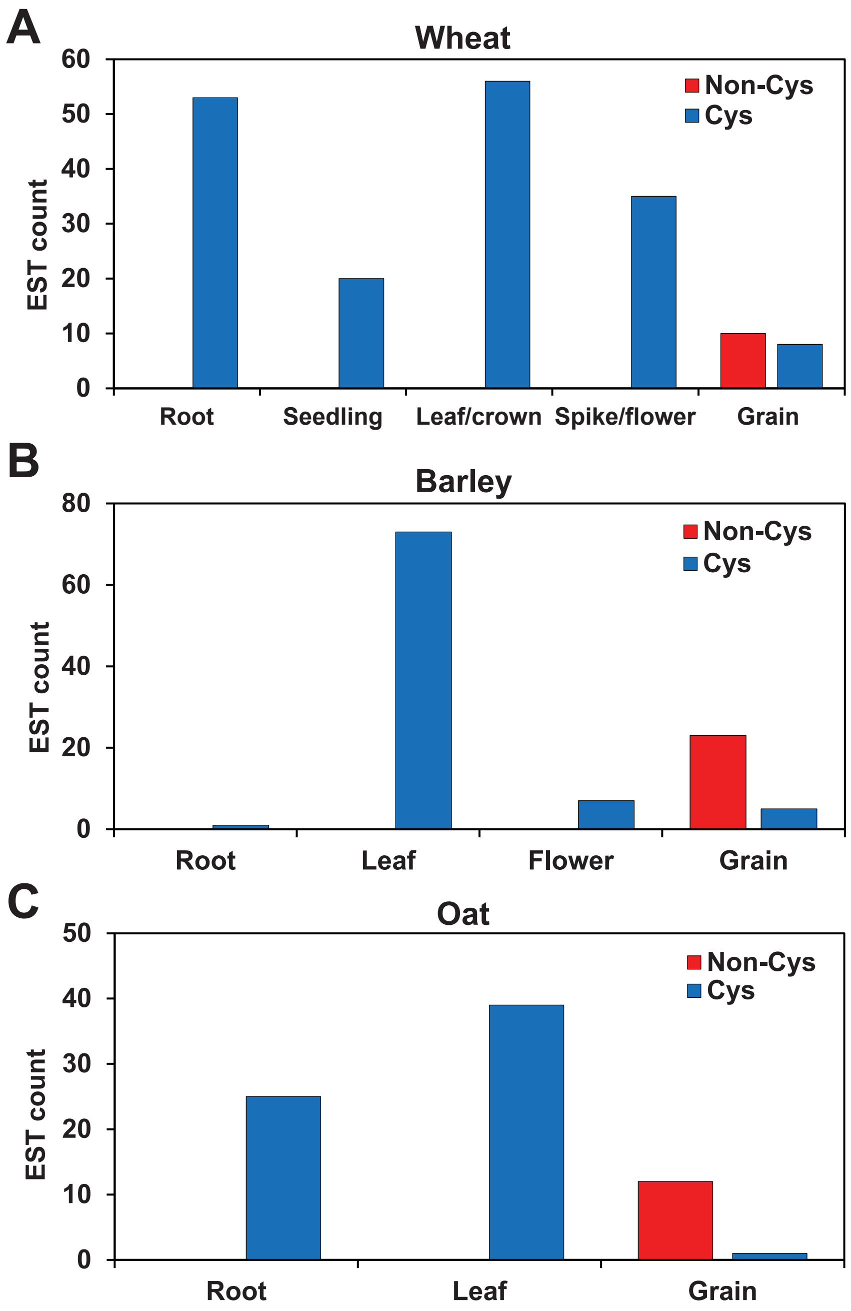


**Supplementary Figure 2. EST evidence that wheat, barley, and oat have canonical Cys THI4s that are widely expressed and non-Cys THI4s that are expressed in developing grains.**
EST counts were taken from the NCBI dbEST database. Only ESTs that could be unambiguously classified as encoding a non-Cys or Cys THI4 and whose source organ was specified were scored.
(**A**) Data for bread wheat. Source cultivars included Chinese Spring, Cheyenne, Cranbrook, Glenlea, Halberd, Jinan 177, Mercia, Recital, Soleil, Sumai3, and Thatcher Lr1.
(**B**) Data for barley. Source cultivars included Barke, Himalaya, Morex, and Optic.
(**C**) Data for oat (*Avena sativa*, cv. CDC Dancer)) and wild oat (*A. barbata*).

```
ScTHI4_3FPZ        1                    MSATSTATSTSASQLHLNSTPVTHCLSDIVKKEDWSDFKFAPIRESTVSRAMTSRYFKDLD
AtThi4_1RP0        1                                              GRGSAGYDLNAFTFDPIKESIVSREMTRRYMTDMI
Barley non-Cys     1                                      MAAICNSISSSTPPYDLNCFKFNPMKESVASREMTRRYMTDMI
Wheat non-Cys      1 MAAMTITTSSFLKTSFPGVCIPAATR--TPSCAATRRTGAICNSISSCTPPYDLSSFKFSPMKESVASREMIRRYMTDMI
Oat non-Cys        1 MAAMATAAPILLKTSFAGACLPKAARSLTLDAVCTPRAGAIYNSISSSTPPYDFNAVKFKPIKESIASREMIRRYMADMI
MiTHI4_4Y4N        1                                 MGSDKIHHHHHHSSGENLYFQGHMDVRLRADEYATTRAILKSAFDMWL

                                   *                    *       *
ScTHI4_3FPZ       62 KFAVSDVIIVGAGSSGLSAAYVIAKNRPDLKVCIIESSVAPGGGSWLGGQLFSAMVMRKPAHLFLQELEIPYED-EGDYV
AtTHI4_1RP0       36 TYAETDVVVVGAGSAGLSAAYEISKN-PNVQVAIIEQSVSPGGGAWLGGQLFSAMIVRKPAHLFLDEIGVAYDE-QDTYV
Barley non-Cys    44 ADVNTDVIIIGTGSAGLSCAYELSKD-PSVNIAIVERSVSPGGSGWLGSQLFSAMVVRKPAHLLLDEINIEYDE-QEDYV
Wheat non-Cys     79 ADVNTDVIIIGTGSAGLSCAYELSKD-PSVNIAIIQRSVSPGGSGWLGSQLFSAMVVRKPAHLFLDELNIEYDE-QEDYV
Oat non-Cys       81 TFSDTDVVIIGTGPAGLSCAYELSKD-PSINIAIIEQSVSPGGSAWLGGQFCSAMVVRKTAHLFLDELNIPYDE-QEDYV
MiTHI4_4Y4N       49 DIIDVDVAIVGGGPSGLTAARYIAKE--GYKVVVLERHLAFGGGTWGGGMGFPYIVVEEPADEILREVGVKLEKVEGEDG

                                                  *                                        *
ScTHI4_3FPZ      141 VVKHAALFISTVLSKVLQLPNVKLFNATCVEDLVTRPPTEKGEVTVAGVVTNWTLVTQAHGT----QCCMDPNVIELAGY
AtTHI4_1RP0      114 VVKHAALFTSTIMSKLLARPNVKLFNAVAAEDLIVK------GNRVGGVVTNWALVAQNHHT----QSCMDPNVMEAK--
Barley non-Cys   122 VIKHAALFTSTLLSRLLAQPNVKLFNAVVVEDLVVK------ENRVAGVITNWALASMNQDI----KSHIDPNVMEGK--
Wheat non-Cys    157 VIKHAALFTSTVLSRLLARPNVKLFNGVVVEDLVVK------EHRVTGVVTNWALVSMNQDTHSQTQSHMDANVMEAK--
Oat non-Cys      159 VIKHAALFTSTILSRLLTCPNVKLFNAVEVEDLIIK------ENHVAGVVTNWALVSMKHDTQ---SYDMDPNIMEAK--
MiTHI4_4Y4N      127 LYTADSVEVPAKLAVGAIDAGAKVLTGIVVEDLVLR------ENRVAGVVINSYAIEKA-------GLHIDPITITAK--

                                         *                                                         *
ScTHI4_3FPZ      217 KNDGTRDLSQKHGVILSTTGHDGPFGAFC--AKRIVDIDQNQKLG--GMKGLDMNHAEHDVVIHSGAYAGVDNMYFAGME
AtTHI4_1RP0      182 -------------IVVSSCGHDGPFGATG--VKRLKSIGMIDHVP--GMKALDMNTAEDAIVRLTREV--VPGMIVTGME
Barley non-Cys   190 -------------IVVSSCGHEGLFSANG--VKRLEDIGTINTMPR--MKALDVNTAEDAIVSLTREV--VPGMIVAGIE
Wheat non-Cys    229 -------------IVVSSCGHEGLFSANGKGVKRLEDIGMIKTVPRTGMEALDTNVSEDAIVGLTREV--VPGMIVAGIE
Oat non-Cys      228 -------------VVVSACGHDRKFGATI--VKHFQD-RMIETMPP-GMSVLDENMSEEDIVHYTREV--VPGLIVTGIQ
MiTHI4_4Y4N      192 -------------YVVDATGHDASVVTTL--SRKNPELGLEVP----GEKSMWAEKGENALLRNTREV--YPGLFVCGMA

                             * *
ScTHI4_3FPZ      293 VAELDGLNRMGPTFGAMALSGVHAAEQILKHFAA
AtTHI4_1RP0      243 VAEIDGAPRMGPTFGAMMISGQKAGQLALKALGLPNAIDGTL
Barley non-Cys   251 VAEIDGPHRMLPTFGATIISGQKAAHLALKALGRPNDIDEQRYSRESRRYTRRLS
Wheat non-Cys    294 VAEIDGPQRMCPTFGATIISGQKAAHLALKALGRPNGIDSETVPA
Oat non-Cys      289 VADIEGAPRMVPTFGATMISGQKAAHLALKALGRPNSIDRTKRCTRSS
MiTHI4_4Y4N      251 ANAVYAGHRMGAIFGGMYISGKKCAEMIVEKLKNNE
```

**Supplementary Figure 3. Alignment non-Cys THI4 sequences from cereals with Arabidopsis and yeast Cys THI4 sequences and the *Methanococcus igneus* non-Cys THI4 sequence.**

Residues in the Arabidopsis (At), yeast (Sc), and *M. igneus* (Mi) THI4s are numbered as in the crystal structures (PDB 1RP0, 3FPZ, 4Y4N, respectively). The active-site Cys in the Arabidopsis and yeast sequences is shaded yellow; residues that replace it in the *M. igneus* and cereal sequences are shaded blue. The $Fe^{2+}$-binding Asp and His residues in the *M. igneus* sequence are shaded red. Active site residues identified in yeast [1,3], Arabidopsis [7], and *M. igneus* [3] THI4 crystal structures are shaded magenta; those identified in all three structures are asterisked. Conserved Cys residues predicted to be in the active site cleft of non-Cys THI4s are shaded cyan. Residue numbers for the full-length Arabidopsis protein can be obtained by adding 44 to the numbers shown. The active-site Cys in yeast THI4 is replaced by dehydroalanine in the crystal structure. The cereal THI4 sequences lack the plastid targeting peptide.

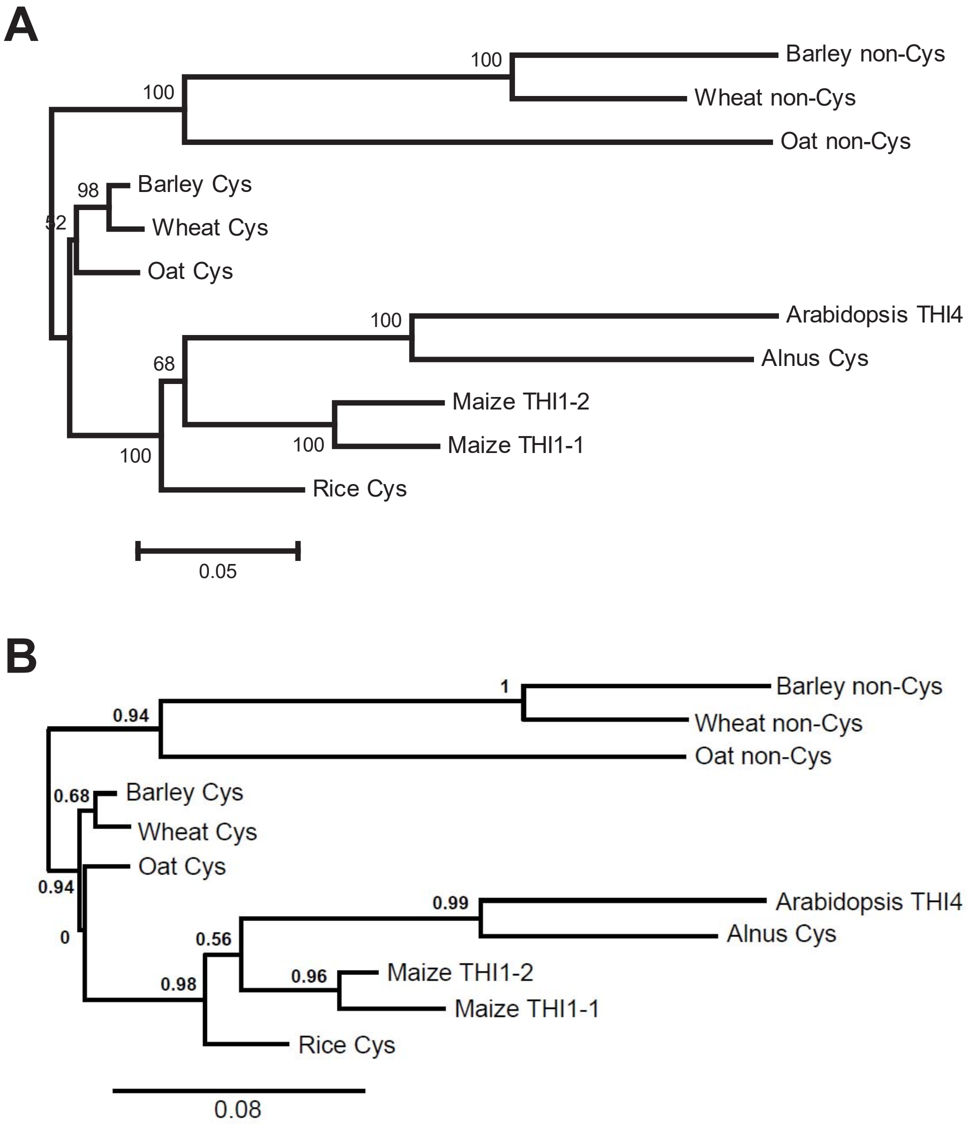


**Supplementary Figure 4. Phylogenetic relationship of cereal non-Cys THI4 sequences to Cys THI4 sequences from monocots and eudicots.**

The THI4 sequences are the same as those in Supplementary Figure 1. Evolutionary distances for each tree are in units of amino acid substitutions per site.

(**A**) Neighbor-joining tree (1,000 bootstrap replicates) built with MEGA 5.2; sequences were aligned with ClustalW. Bootstrap values are shown for branch support.

(**B**) Tree built with Phylogeny.fr; sequences were aligned with Muscle. Approximate likelihood ratios are shown for branch support.

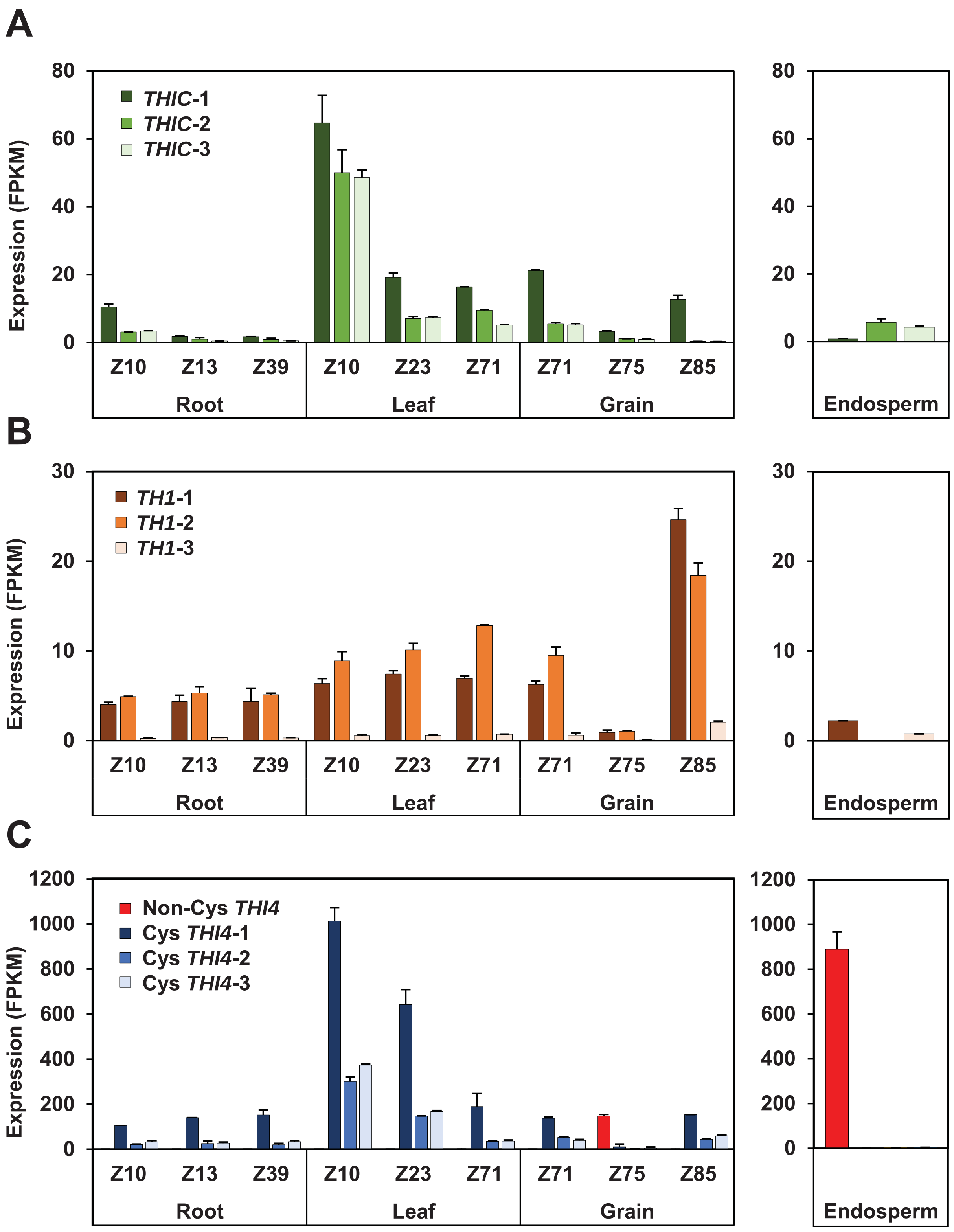


**Supplementary Figure 5. RNA-seq evidence for the expression of wheat thiamin synthesis enzymes THIC and TH1 in developing grains.**

Expression of the three *THIC* (**A**), three *TH1* (**B**), and three Cys *THI4* genes and the single non-Cys *THI4* gene (**C**) of wheat cv. Chinese Spring in roots, leaves, and grains at various stages on the Zadoks (Z) growth stage scale, and in endosperm. Values are means and s.e.m. of two replicates. Data are from https://wheat.pw.usda.gov/WheatExp/. Gene identifiers are listed in Supplementary Table 3.

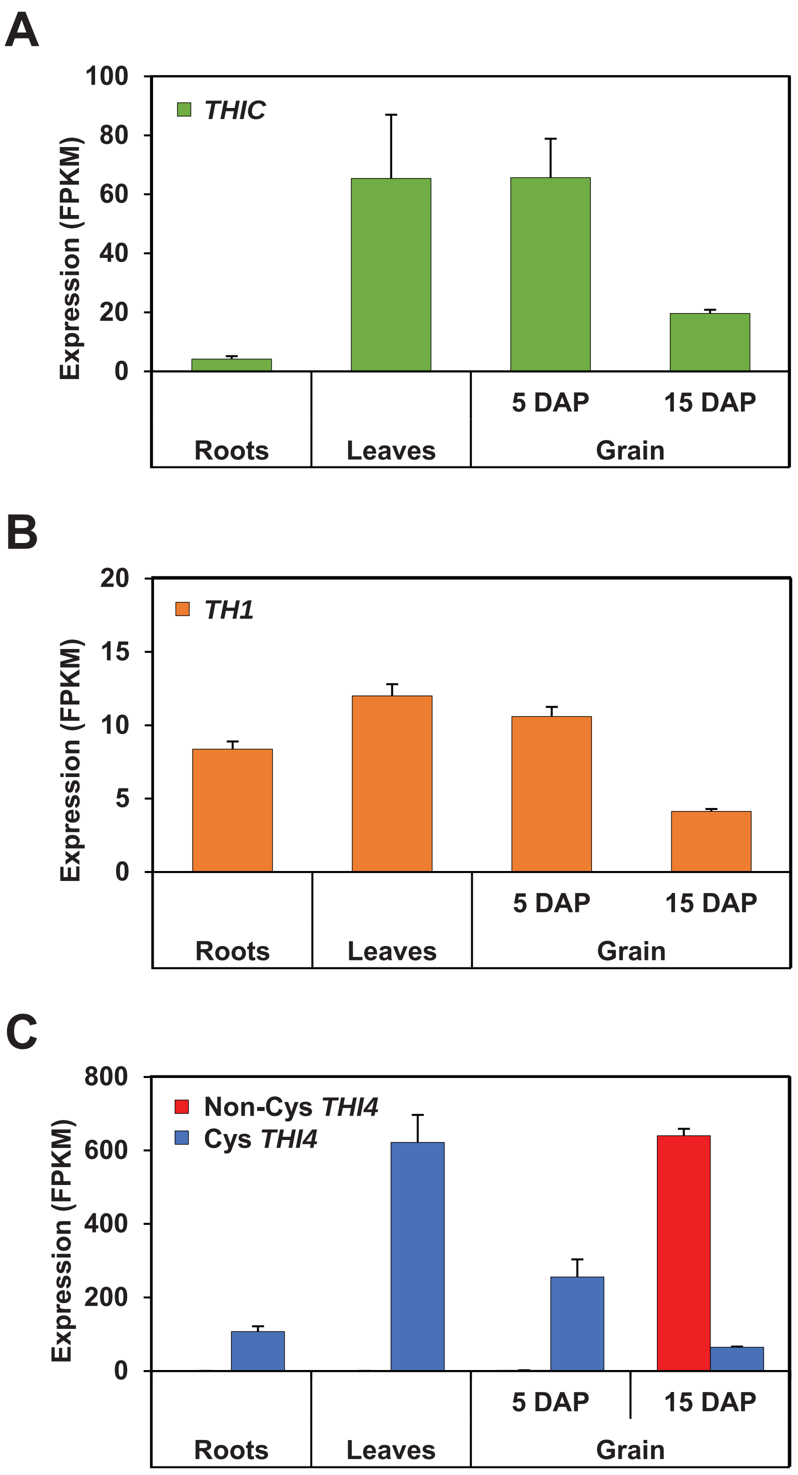


**Supplementary Figure 6. RNA-seq evidence for the expression of barley thiamin synthesis enzymes THIC and TH1 in developing grains.**

Expression of the single *THIC* (**A**), *TH1* (**B**), and non-Cys and Cys *THI4* genes (**C**) of barley cv. Morex in roots, leaves, and grains at two developmental stages. DAP, days after pollination. Values are means and s.e.m. of 3 replicates. Data are from https://ics.hutton.ac.uk/morexGenes/index.html. Gene identifiers are listed in Supplementary Table 3.

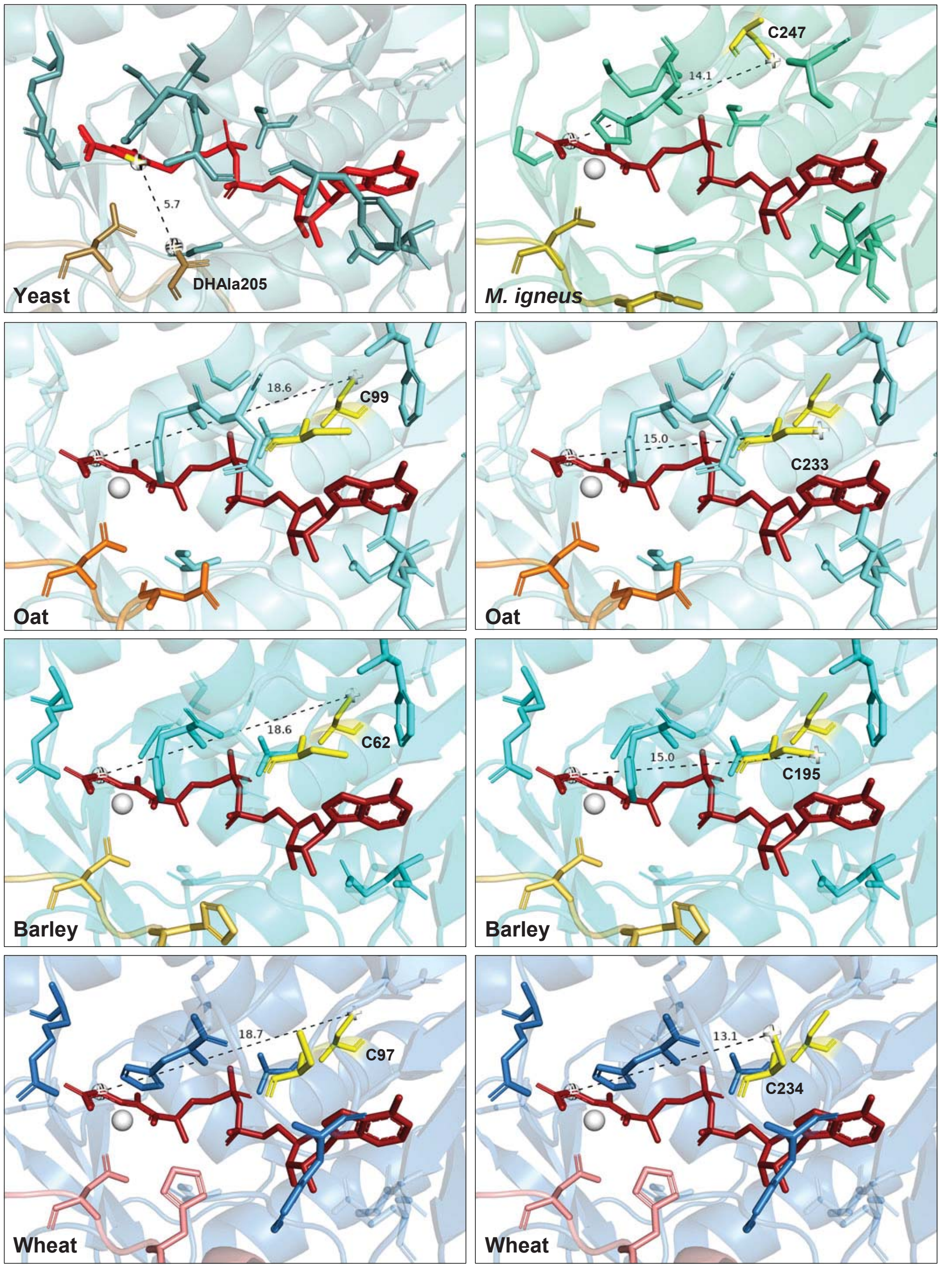


**Supplementary Figure 7. Cys residues in non-Cys THI4s that are unlikely to donate sulfur.**

Top left frame: As a benchmark, the DHAla 205 (formerly C205) residue of yeast Cys THI4 (PDB 3FPZ) is 5.7 Å from the sulfur atom (yellow) in the bound ADP-5-ethyl-4-methylthiazole-2-carboxylate reaction product (bright red).

Other frames: The single active site cleft Cys residue of *M. igneus* THI4 (PDB 4Y4N) and both active site cleft Cys residues of cereal non-Cys THI4s are located >13 Å from the substrate, glycine imine (brick red). For cereal THI4s, the left-hand column shows the Cys residue in the residue 62-99 region; the right-hand column shows the Cys residue in the residue 195-234 region. Glycine imine and iron (white) were modeled into the active sites using the Pymol 2.3.5 alignment command (template: 4Y4N).

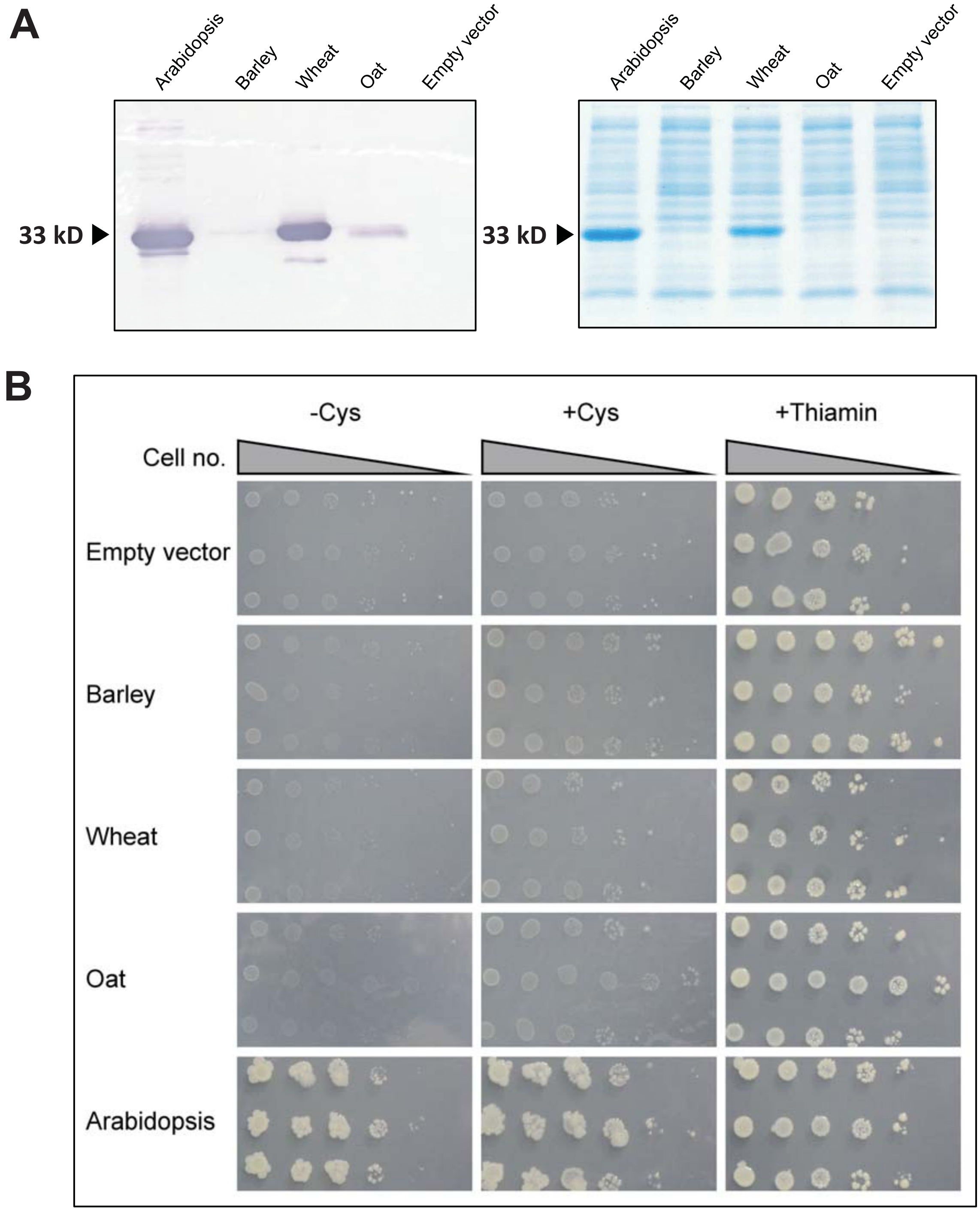


**Supplementary Figure 8. Expression levels and complementing activity of non-Cys THI4s in *E. coli*.**
(**A**) Expression of $His_6$-tagged barley, wheat, and oat non-Cys THI4s in strain BL21(DE3). The Cys THI4 of Arabidopsis and an empty pET28b vector control are also shown. Lanes contained 5 µg of soluble protein extract. THI4 proteins were detected by Western blotting using anti-$His_6$ antibodies (left) or Coomassie blue staining (right).
(**B**) Complementation tests in a BL21(DE3) Δ*thiG* strain. Cells harboring the pET28b vector alone or containing a $His_6$-tagged THI4 were cultured in MOPS medium supplemented with 0.2% glucose and 1 mm IPTG, minus or plus 1 mM Cys, in aerobic conditions. Controls supplemented with 100 nM thiamin were included. Overnight liquid cultures of three independent clones of each construct were serially diluted in ten-fold steps and spotted on the plates. Images were captured after incubation at 37°C for 6 d (empty vector and cereal THI4s) or 2 d (Arabidopsis THI4).

**Supplementary Table 1 Oligonucleotide primers used in this study**

| Primer | Name | Sequence 5' → 3' |
|---|---|---|
| | | **AtTHI4 promoter/coding sequence** |
| 1 | AtTHI4_promoter | GGTACCGCTATCTAGTGGATCATATTATTAATAAAACC |
| 2 | | GGATCCAGTCGCTAGCAGCATCTACGGTTTCAGCTG |
| 3 | AtTHI4_3´ sequence | AGTCGCTAGCCATCATCACCATCACCATTGAAATCAAAAG GTTTTAACGTTAAGAAAAG |
| 4 | | AGTCGGATCCCATTGCAGAAGATTCACGAG |
| | | **Construction of *E. coli* BL21(DE3) Δ*thiG*** |
| 5 | 5' ThiG | ATATCGTGCAGGATGGC |
| 6 | 3' ThiG | CGTTGATACGCAGGCGG |
| 7 | ThiG upstream | CGACCAACGACAAGCG |
| 8 | Kan 3' | GATGCGCTGCGAATCGGG |
| 9 | ThiG 5' | GCCAGCACAACGACGC |
| | | **pET28b-Cereal THI4** |
| 11 | 5'AtThi4Δ47XbaI | CGTATCTAGAAATAATTTTGTTTAACTTTAAGAAGGAGATAT ACCATGACCCCGCCGTATGACCTGAATTGC |
| 12 | 3'AtThi4Δ47XhoI | CGTACTCGAGAGCATCTACGGTTTCAGCTGAATCTGCTGC |
| 13 | 5'HvThi4Δ46XbaI | CGTATCTAGAAATAATTTTGTTTAACTTTAAGAAGGAGATAT ACCATGACCCCGCCGTATGACCTGAATTGC |
| 14 | 3'HvThi4Δ46XhoI | CGTACTCGAGGCTCAGACGGCGGGTGTAGC |
| 15 | 5'TaThi4Δ46XbaI | CGTATCTAGAAATAATTTTGTTTAACTTTAAGAAGGAGATAT ACCATGACCCCGCCGTACGATCTGAGC |
| 16 | 3'TaThi4Δ46XhoI | CGTACTCGAGCGCCGGAACGGTTTCGCTATCAATACC |
| 17 | 5'AsThi4Δ48XbaI | CGTATCTAGAAATAATTTTGTTTAACTTTAAGAAGGAGATAT ACCATGACCCCGCCGTATGATTTCAACG |
| 18 | 3'AsThi4Δ48XhoI | CGTACTCGAGGCTACGGGTGCAACGCTTGGTACG |

**Supplementary Table 3 Gene identifiers for wheat and barley *THI4* transcripts**

| | Legend ID | Transcript ID |
|---|---|---|
| Wheat | Non-Cys THI4 | TRAES3BF087900030CFD |
| | Cys THI4-1 | Traes_7DL_51CA70B80.1 |
| | Cys THI4-2 | Traes_7AL_6BDC0C3EA.1 |
| | Cys THI4-3 | Traes_7AL_4C91CAF32.1 |
| | THIC-1 | Traes_4DS_0A7E021B3.2 |
| | THIC-2 | Traes_4BS_E79CB87B7.2 |
| | THIC-3 | Traes_4AL_2A2E81E05.5 |
| | TH1-1 | Traes_6BS_655F3A82D.1 |
| | TH1-2 | Traes_6AS_1EF252E0A.1 |
| | TH1-3 | Traes_6DS_7BDE4D0B8.1 |
| Barley | Non-Cys THI4 | MLOC_5785 |
| | Cys THI4 | MLOC_11312 |
| | THIC | MLOC_43949 |
| | TH1 | MLOC_59029 |

**Supplementary Table 4  Nucleotide sequences of recoded cereal THI4s**

The barley sequence lacks part of the targeting peptide; the wheat and oat sequences are full length.

| Species | Recoded sequences (5´→ 3´) |
|---|---|
| Barley | CC**ATG**GCGGCGATTTGCAACAGCATTAGCAGCAGCACCCCGCCGTATGACCTGAATTGCT TTAAGTTTAACCCGATGAAGGAGAGCGTGGCGAGCCGTGAGATGACCCGTCGTTACATGA CCGACATGATTGCGGATGTTAACACCGACGTGATCATTATTGGTACCGGCAGCGCGGGTC TGAGCTGCGCGTATGAGCTGAGCAAGGACCCGAGCGTTAACATTGCGATCGTTGAACGTA GCGTGAGCCCGGGTGGCAGCGGTTGGCTGGGCAGCCAGCTGTTCAGCGCGATGGTGGT TCGTAAGCCGGCGCACCTGCTGCTGGACGAGATTAACATCGAATACGATGAGCAGGAAGA CTATGTGGTTATCAAACACGCGGCGCTGTTCACCAGCACCCTGCTGAGCCGTCTGCTGGC GCAACCGAACGTGAAGCTGTTTAACGCGGTGGTTGTGGAGGATCTGGTTGTGAAAGAAAA CCGTGTTGCGGGTGTGATTACCAACTGGGCGCTGGCGAGCATGAACCAAGATATTAAGAG CCACATCGACCCGAACGTTATGGAGGGCAAAATCGTTGTGAGCAGCTGCGGTCACGAAGG CCTGTTCAGCGCGAACGGTGTGAAGCGTCTGGAAGATATTGGCACCATCAACACCATGCC GCGTATGAAAGCGCTGGATGTTAACACCGCGGAGGACGCGATTGTGAGCCTGACCCGTG AAGTTGTGCCGGGTATGATTGTTGCGGGCATCGAGGTGGCGGAAATCGACGGTCCGCAC CGTATGCTGCCGACCTTTGGTGCGACCATTATTAGCGGTCAGAAGGCGGCGCACCTGGC GCTGAAAGCGCTGGGCCGTCCGAACGACATTGATGAGCAACGCTACAGCCGTGAGAGCC GCCGCTACACCCGCCGTCTGAGC**TGA**TCTAGA |
| Wheat | GTAAAACGACGGCCAGTGCC**ATG**GCGGCGATGACCATTACCACCAGCAGCTTTCTGAAGA CCAGCTTTCCGGGCGTGTGCATTCCGGCGGCGACCCGTACCCCGAGCTGCGCGGCGACC CGTCGTACCGGTGCGATTTGCAACAGCATCAGCAGCTGCACCCCGCCGTACGATCTGAGC AGCTTCAAGTTTAGCCCGATGAAAGAGAGCGTTGCGAGCCGTGAAATGATTCGTCGTTATA TGACCGACATGATCGCGGATGTTAACACCGACGTGATCATTATTGGTACCGGTAGCGCGG GTCTGAGCTGCGCGTACGAGCTGAGCAAGGATCCGAGCGTTAACATTGCGATTATTCAAC GTAGCGTGAGCCCGGGTGGCAGCGGTTGGCTGGGCAGCCAACTGTTCAGCGCGATGGTG GTTCGTAAGCCGGCGCACCTGTTTCTGGACGAGCTGAACATTGAATACGATGAGCAGGAA GACTATGTGGTTATCAAACACGCGGCGCTGTTCACCAGCACCGTTCTGAGCCGTCTGCTG GCGCGTCCGAACGTGAAGCTGTTTAACGGTGTGGTTGTGGAGGACCTGGTTGTGAAAGAA CACCGTGTTACCGGCGTTGTGACCAACTGGGCGCTGGTGAGCATGAACCAAGATACCCAC AGCCAGACCCAAAGCCACATGGACGCGAACGTGATGGAGGCGAAGATCGTTGTGAGCAG CTGCGGTCACGAAGGCCTGTTCAGCGCGAACGGCAAGGGCGTTAAACGTCTGGAGGATA TTGGTATGATCAAAACCGTGCCGCGTACCGGTATGGAGGCGCTGGATACCAACGTTAGCG AAGACGCGATTGTGGGTCTGACCCGTGAAGTTGTGCCGGGTATGATTGTTGCGGGCATTG AGGTGGCGGAAATTGATGGTCCGCAGCGTATGTGCCCGACCTTTGGTGCGACCATTATCA GCGGCCAAAAAGCGGCGCACCTGGCGCTGAAGGCGCTGGGTCGTCCGAATGGTATTGAT AGCGAAACCGTTCCGGCG**TAA**TCTAGACATGGTCATAGCTGTTTCC |
| Oat | TAATACGACTCACTATAGGGCC**ATG**GCGGCGATGGCGACCGCGGCGCCGATCCTGCTGA AGACCAGCTTTGCGGGTGCGTGCCTGCCGAAGGCGGCGCGTAGCCTGACCCTGGACGC GGTTTGCACCCCGCGTGCGGGTGCGATCTACAACAGCATCAGCAGCAGCACCCCGCCGT ATGATTTCAACGCGGTGAAGTTTAAACCGATCAAGGAGAGCATTGCGAGCCGTGAAATGAT CCGTCGTTACATGGCGGACATGATTACCTTTAGCGACACCGATGTGGTTATCATTGGTACC GGTCCGGCGGGTCTGAGCTGCGCGTATGAGCTGAGCAAAGATCCGAGCATCAACATTGC GATCATTGAACAGAGCGTTAGCCCGGGTGGCAGCGCGTGGCTGGGTGGCCAATTCTGCA GCGCGATGGTGGTTCGTAAGACCGCGCACCTGTTTCTGGACGAGCTGAACATCCCGTACG ACGAGCAGGAAGATTATGTGGTTATCAAACACGCGGCGCTGTTCACCAGCACCATTCTGA GCCGTCTGCTGACCTGCCCGAACGTGAAGCTGTTTAACGCGGTTGAGGTGGAAGACCTGA TCATTAAAGAGAACCACGTTGCGGGTGTGGTTACCAACTGGGCGCTGGTTAGCATGAAGC ACGATACCCAGAGCTACGACATGGATCCGAACATCATGGAAGCGAAAGTGGTTGTGAGCG CGTGCGGTCACGACCGTAAGTTCGGCGCGACCATCGTTAAACACTTTCAAGATCGTATGAT TGAGACCATGCCGCCAGGCATGAGCGTTCTGGACGAAAACATGAGCGAGGAAGATATTGT GCACTATACCCGTGAGGTTGTGCCGGGTCTGATCGTTACCGGCATTCAGGTGGCGGACAT CGAAGGTGCGCCGCGTATGGTGCCGACCTTTGGTGCGACCATGATTAGCGGTCAAAAAGC GGCGCACCTGGCGCTGAAAGCGCTGGGCCGTCCGAACAGCATCGATCGTACCAAGCGTT GCACCCGTAGCTC**TAG**ACTATAGTGTCACCTAAATC |

**Supplementary Table 5  RNA-seq data showing the predominant expression of a non-Cys THI4 gene in developing seeds of oat genotype Ogle-C**

Four THI4 genes (loci) were detectably expressed in developing seeds harvested at 7, 14, 21, and 28 days after anthesis, and pooled for RNA preparation [18]. Data are mean and s.e.m. for hits on two to five transcript models per locus.

| Locus | THI4 type | RPKM |
|---|---|---|
| 84 | Non-Cys | 419 ± 43 |
| 2457 | Cys | 42 ± 1 |
| 3337 | Cys | 84 ± 2 |
| 12043 | Cys | 21 ± 0.4 |

**Supplementary Table 6  Residue differences between cereal non-Cys THI4s and other THI4s**

Residue conservation is based on searches of the ~500 Cys THI4 sequences from angiosperms in the NCBI nr protein database and of 114 prokaryote Cys and non-Cys THI4 sequences curated in the SEED database. Residues are numbered using the full length Arabidopsis THI4 sequence. $Cys^{216}$ is the active-site Cys residue. In addition to the tabulated single-residue differences, cereal THI4s have small insertions after $Thr^{213}$ and $Pro^{255}$. The residue numbers in the Arabidopsis THI4 crystal structure [7] can be obtained by subtracting 44 from the numbers in the table.

| Arabidopsis THI4 residue | Conserved in angiosperms | Conserved in prokaryotes | Cereal non-Cys THI4 residues |
|---|---|---|---|
| $Val^{66}$ | Yes | No | Ala |
| $Ala^{91}$ | Yes | No | Thr |
| $Gly^{122}$ | Yes | Yes | Ser |
| $Cys^{216}$ | Yes | No | His or Asp |
| $Pro^{236}$ | Yes | No | Leu or Lys |
| $Gly^{297}$ | Yes | Yes | Leu, Cys, or Val |
| $Met^{303}$ | Yes | Yes | Thr |
